# Smad7-based biologic targeting epidermis and stroma promotes healing of diabetic wounds in mice and pigs

**DOI:** 10.64898/2025.12.31.697220

**Authors:** Yao Ke, Ben-Zheng Li, Fulun Li, Resmi K. Ravindran, Donna Wang, Suyan Wang, Sam T Hwang, Scott Simon, Sean R. Collins, Christian D. Young, Xiao-Jing Wang

## Abstract

This study aimed to identify novel mechanisms of diabetic wound healing defects and test a therapeutic intervention using diabetic mouse and pig models. We found Smad7 transgene expression in mouse epidermis promoted wound healing in diabetic mice. To isolate effects of Smad7 on wounds, we created a Smad7-based biologic (Tat-PYC-Smad7) that penetrated cells of the wound. Topical Tat-PYC-Smad7 treatment to diabetic pig and mouse wounds accelerated healing compared to vehicle controls. Tat-PYC-Smad7-treated wounds showed reduced TGFβ/NFκB signaling, faster re-epithelialization and better extracellular matrix remodeling. Tat-PYC-Smad7 also attenuated neutrophil NETosis, potentially acting through reductions in MPO enzymatic activity and MPO nuclear entry, consequently reducing chromatin decondensation and the release of NET components. Our study revealed that Tat-PYC-Smad7 promoted diabetic wound healing by targeting keratinocytes and neutrophils, providing insight into mechanisms of diabetic wound healing defects targetable by Smad7-based therapy.

## Introduction

Impaired wound healing in diabetes represents a significant cause of morbidity ^1^. Standard wound management is often ineffective in treating diabetic wounds, posing global health, economic, and social issues ^2, 3^. Becaplermin/Regranex, a recombinant platelet-derived growth factor (PDGF)-BB, is the only FDA-approved topical drug for diabetic foot ulcers ^4^. However, Regranex requires large, repetitive doses to achieve even modest improvement in a small proportion of patients ^5^. In addition, long-term use of growth factors is linked to a carcinogenesis risk ^6^. Hence, developing novel, effective treatments for diabetic wounds is an unmet medical need.

The wound healing process includes stages of inflammation, proliferation, and remodeling ^3^. In diabetic wounds, healing stalls at the inflammation phase ^7^. One of the hallmarks of nonhealing human wounds is keratinocyte dysfunction ^8^, which has been considered a therapeutic target for diabetic wounds^9^. Prolonged inflammation in nonhealing wounds generates reactive oxygen species (ROS), leading to keratinocyte injury, apoptosis, and impaired migration ^10^. Moreover, altered extracellular matrix (ECM) composition and enhanced ECM degradation by elevated matrix metalloproteinase levels in nonhealing wounds impair keratinocyte attachment, leading to aberrant cell signaling and impaired survival and migration ^11^. In addition to keratinocyte dysfunction, prolonged infiltration and activation of inflammatory cells, mainly neutrophils, are the main source of proinflammatory cytokines, proteases, and ROS in chronic wounds ^12^. This contributes to the recruitment of more immune cells, alterations in the proteolytic balance, and impaired blood vessel formation ^13^. These dysregulations delay wound healing, exacerbate scar formation, and predispose local lesions to neoplastic progression ^7, 14^. In patients with diabetes, elevated glucose levels promote neutrophil extracellular trap (NET) formation through the process of NETosis ^15^. The pathogenic role of prolonged neutrophil activity in diabetic wounds potentially serves as a therapeutic target. Conversely, individuals with neutropenia or those suffering from conditions with neutrophil abnormalities (such as deficiencies in adhesion or neutrophil granules etc.) are more susceptible to wound infections resulting in abnormal wound healing processes ^16^. This suggests that a systemic approach to targeting neutrophils may not be viable because neutrophils play dual roles in wound healing.

To study mechanisms of healing defects in diabetic wounds and test therapeutic interventions, animal models have been developed ^17^. Excisional wounds in diabetic BKS.Cg-Dock7m +/+ Leprdb/J (db/db) mice have a significant healing delay, with reductions in neovascularization and epidermal proliferation ^18^. Miniature diabetic pigs are also a model for wound healing studies due to the similarities of their skin texture to humans ^19^. While the porcine model offers a clinically relevant model, e.g. enabling an assessment of treatment effects on ECM remodeling, its limitations, such as the extensive clinical care, long preparation time, and limited excisional wounds per pig restrict its feasibility for mechanistic investigations. Therefore, mouse models provide a complementary resource for mechanistic studies. We previously observed that transgenic mice constitutively overexpressing wild-type TGFβ1 in keratinocytes (K5.TGFβ1^wt^) with TGFβ/NFκB signaling activation exhibited a significant delay in full-thickness wound healing, characterized by profound inflammation, keratinocyte apoptosis, and ECM dysregulation ^20, 21^. We also found that TGFβ and NFκB are activated in human diabetic wounds ^21^, suggesting that a TGFβ/NFκB dual inhibitor, Smad7, may have therapeutic effects on diabetic wounds. Smad7 consists of an N-terminal and C-terminal domain, with a linker region containing a PY motif ^22^. The functional activities of Smad7, including blocking TGFβ/NFκB signaling, are attributed to its C-terminal domain and PY motif ^23^. These domains of human Smad7 share 99% amino acid homology with mouse and pig Smad7, enabling the use of these animal models to investigate its function while minimizing species-mismatch-associated immunogenic responses ^22^. In this study, we employed both genetic and pharmaceutical models to assess the role of Smad7 in diabetic wound repair. We provide evidence that Smad7-based proteins promoted diabetic wound healing via supporting epithelial cell survival, ECM modification, and attenuating neutrophil activities.

## Results

### K5.Smad7 transgene expression and Tat-PYC-Smad7 topical application mitigated wound healing defects of db/db mice

We previously developed K5.Smad7 mice, which express Smad7 in keratinocytes ^24^. We bred K5.Smad7 mice with db/db mice that have obesity and type 2 diabetes ^25^, to generate K5.Smad7/db/db bigenic mice. Grossly, K5.Smad7/db/db mice had sparse hair (Fig.1A), similar to K5.Smad7 mice ^26^. The db/db phenotype of obesity and high blood glucose levels were significantly lower in K5.Smad7/db/db mice than in db/db littermates (Fig.1B-C, Sup Table 1). Microscopically, K5.Smad7/db/db skin showed mild epidermal hyperplasia and hypertrophic sebaceous glands, similar to K5.Smad7 skin (Fig.1A). To assess whether the K5.Smad7 transgene could attenuate wound healing defects of db/db mice, we established excisional wounds by skin punch (6mm) on the backs of mice and observed wound healing kinetics. Compared to db/db mice, wound healing in K5.Smad7/db/db mice was significantly accelerated from day 9 (Fig.1D-E).

**Fig. 1.**
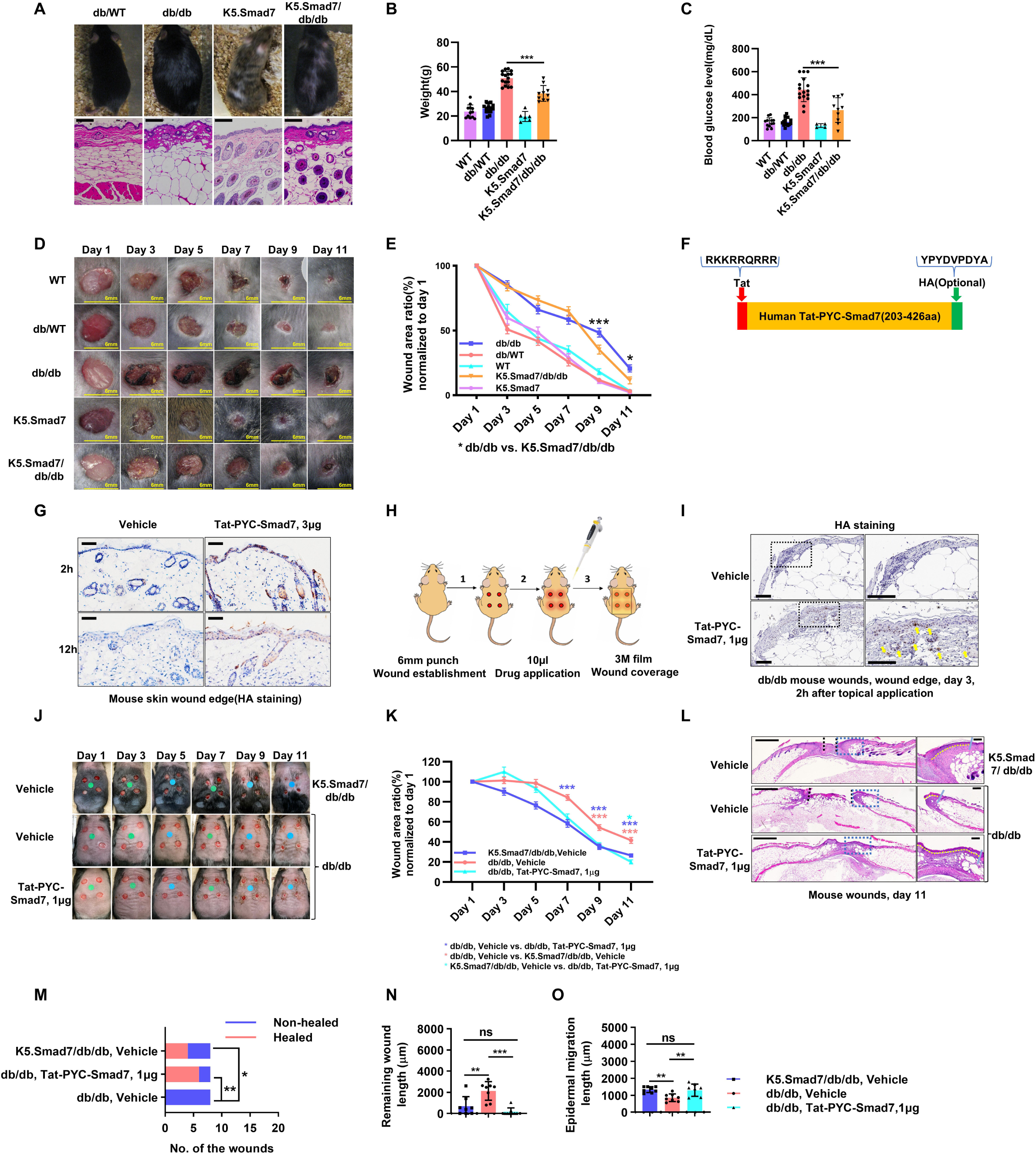
K5.Smad7 transgene and Tat-PYC-Smad7 mitigated wound healing defects in db/db mice. (**A**) Gross photos show the skin phenotype and body size of female K5.Smad7/db/db mice and their littermates with representative H&E images of skin sections. (**B**) Body weight and (**C**) blood glucose levels at the beginning of wound experiments are shown (mean ± SD, n = 8-12/group, both sexes). (**D**) Representative wound photos and (**E**) wound area ratios are shown (mean ± SEM, n = 6-15/group, pooled from two independent experiments). (**F**) A schematic of Tat-PYC-Smad7 protein structure. (**G**) HA immunohistochemistry demonstrates Tat-PYC-Smad7 localization at wound edges 2 and 12 h after topical application. (**H**) Scheme for wound establishment and topical treatment. (**I**) HA staining of db/db wounds 2 h after Tat-PYC-Smad7 or vehicle application. Yellow arrows indicate Tat-PYC-Smad7 nuclear localization. (**J**) Representative wound images(Colored dot stickers with 6 mm in diameter were placed at the center of each wound served as a scale reference) and (**K**) wound area ratios (mean ± SEM, n = 7-10/group, pooled from two independent experiments). (**L**) Representative H&E images of wounds on day 11. Dotted yellow lines mark the migrating epidermal tongue and blue lines mark the leading edge. (**M**) Quantification of healed versus non-healed wounds (n = 8-16/group). (**N**) Quantification of remaining wound length (mean ± SD, n = 8/group). (**O**) Quantification of epidermal migration length (mean ± SD, n = 8/group). Scale bars: 100 µm (**A, G, I**); 2 mm (left) and 200 µm (right) (**M**). \**P* < 0.05, \*\**P* < 0.01, \*\*\**P* < 0.001. WT, wild type.

As K5.Smad7 mice expressed Smad7 transgene in other epithelial cells in addition to keratinocytes ^24^, to assess whether Smad7 healing is a direct effect on wounds or a secondary effect of attenuated diabetic phenotype or from other tissues, we generated a Tat-PYC-Smad7 protein that contains the functional domains of human Smad7 with a cell-penetrating Tat peptide. An optional HA tag was added to the C-terminus of Tat-PYC-Smad7 enabling tracking of this exogenous protein (Fig.1F). Testing topical application of Tat-PYC-Smad7 on normal mouse skin wounds indicated a relatively long stability of Tat-PYC-Smad7 (at least 12h) (Fig.1G). Four full-thickness skin wounds were induced using a skin punch on the dorsum of mice on day 1, followed by topical treatment every other day and application of 3M film after treatment to prevent scab formation (Fig.1H). Tat-PYC-Smad7 penetrated both epithelial and stromal cells of db/db wounds (Fig.1I). Grossly, we observed accelerated wound healing in both K5.Smad7/db/db wounds and wounds of db/db mice treated with Tat-PYC-Smad7 from day 7 to day 11 (Fig.1J-K). Histologically, K5.Smad7 transgene expression or Tat-PYC-Smad7 treatment led to increased diabetic wound healing rates (Fig.1L-M). Additionally, there was a reduction in the remaining wound length and an increase in the length of the epidermal migrating tongue in K5.Smad7/db/db wounds and Tat-PYC-Smad7-treated wounds, compared to vehicle-treated db/db wounds (Fig.1L, N, O). Notably, Tat-PYC-Smad7 topical application exerted an effect comparable to K5.Smad7 transgene expression in promoting diabetic wound healing. We did not observe a significant change of either blood glucose or body weight in mouse hosts with Tat-PYC-Smad7-treated wounds compared to the vehicle control (Sup Fig.1).

We found a dose-dependent effect of Tat-PYC-Smad7 in accelerating wound healing (Sup Fig.2). Then we used 1µg of Tat-PYC-Smad7, which showed the best treatment efficacy, to compare with standard-of-care treatment, Regranex (Sup Fig.3A). We found that Tat-PYC-Smad7 significantly expedited wound closure compared to either vehicle or Regranex treatment on day 5 and 7 (and comparable healing to Regranex at later time points) and Tat-PYC-Smad7 had an overall higher rate of healing (Sup Fig.3B-E). From day 8 to day 11, wounds treated with Tat-PYC-Smad7 exhibited reduced size and longer lengths of epidermal migrating tongues compared to vehicle (similar or better than Regranex-treated wounds) (Sup Fig.3F-H). We also observed foam-like F4/80+ myeloid cells in Regranex-treated wounds but not in Tat-PYC-Smad7-treated wounds (Sup Fig.4). These foam-like cells with internalized lipids and a characteristic foamy cytoplasmic appearance may indicate active inflammation and have been linked to a profibrotic phenotype ^27^. Regranex-treated mice had increased numbers of circulating white blood cells including lymphocytes, monocytes, and granulocytes compared to vehicle- or Tat-PYC-Smad7 treated mice (Sup Fig.5A). In addition, we found that Regranex, but not Tat-PYC-Smad7, upregulated pro-angiogenic factor Angiopoietin 2, chemokines CXCL16 and CXCL5, LDL receptor (LDL R) and IGFBP3 (Sup Fig.5B and C). Angiopoietin 2 is a pro-angiogenic factor that promotes vascular leakage, which has been involved in chronic inflammation and fibrosis ^28^. Chemokines such as CXCL16 and CXCL5 recruit immune cells to inflamed tissues, facilitating fibrotic processes ^29^. LDL R increases oxidized low-density lipoprotein accumulation, activating macrophages and endothelial cells to produce pro-inflammatory cytokines and reactive oxygen species, promoting fibrosis ^30^.

**Fig. 2.**
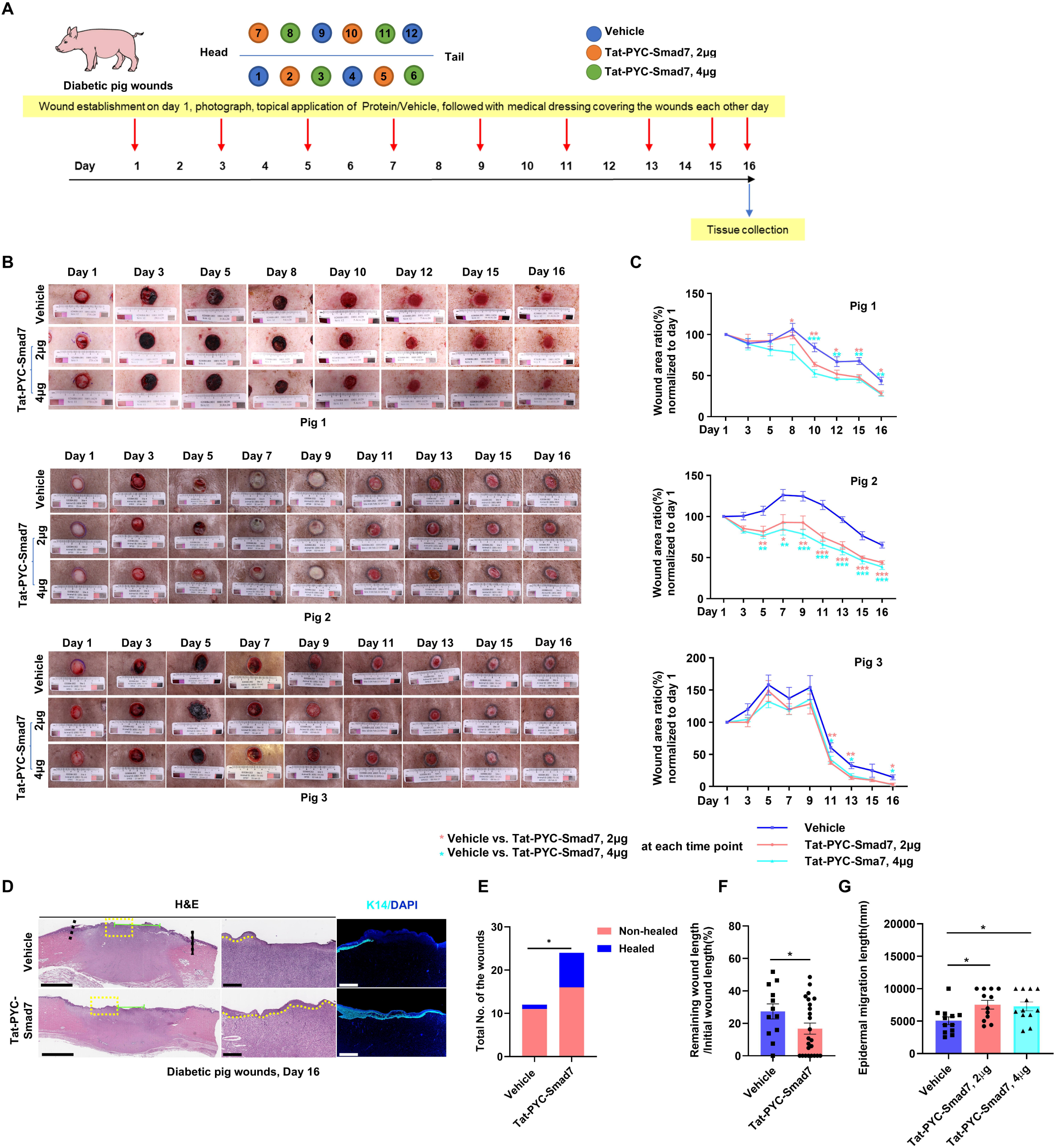
Tat-PYC-Smad7 promoted wound healing in diabetic pigs. (**A**) Schematic for 2-cm full-thickness wounding and Tat-PYC-Smad7 treatment in diabetic pigs. (**B**) Representative gross images and (**C**) quantification of the wound area for each of the three pigs (mean ± SEM, 12 wound samples per pig). (**D**) Representative H&E and immunofluorescent staining of K14 and DAPI in skin wound sections collected on day 16, with the area enclosed by the dotted yellow box magnified and shown on the right. Dotted yellow lines indicate the migrating tongue and dotted black lines indicate the edge of the wound. Scale bars: 4 mm (left), 1 mm (right). **(E)** Wound healing rate of the three pigs, and **(F)** quantification of the remaining wound length normalized to the initial wound length of the three pigs (mean ± SD, n = 12-24/group). **(G)** Quantification of the epidermal migration length of the three pigs by each treatment group(n = 12, mean ± SD). \**P* < 0.05, \*\**P* < 0.01, \*\*\**P* < 0.001. H&E, hematoxylin and eosin; K14, keratin 14.

**Fig. 3.**
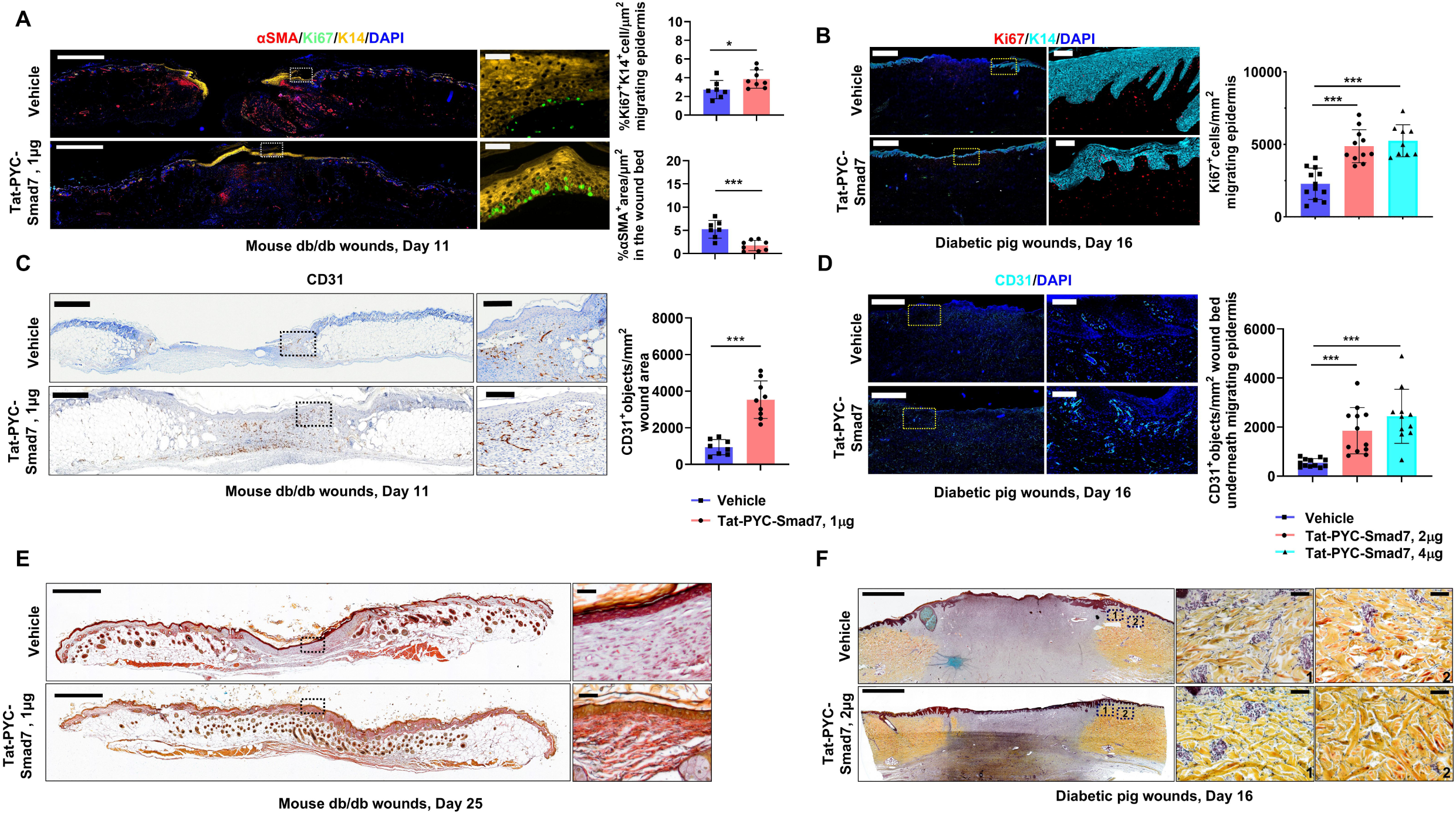
Histological analysis revealed beneficial alteration of wound healing kinetics with Tat-PYC-Smad7 treatment. (**A**) Representative immunofluorescent images of mouse db/db wounds on day 11 stained for αSMA, Ki67, K14, and DAPI, with corresponding quantification of αSMA+ area in the wound bed and Ki67^+^K14^+^ cells in the migrating epidermis (mean ± SD, n = 7-8/group). (**B**) Representative immunofluorescent images of diabetic pig wounds on day 16 stained for Ki67, K14, and DAPI, with quantification of Ki67+ cells in the migrating epidermis (mean ± SD, n = 8-12/group). (**C**) Representative immunohistochemistry images of mouse db/db wounds on day 11 stained for CD31 with quantification of CD31^+^ cells per mm² wound area (mean ± SD, n = 8-9/ group). (**D**) Representative immunofluorescent images of diabetic pig wounds on day 16 stained for CD31 and DAPI with quantification of CD31^+^ cells in the wound bed underneath the migrating epidermis (mean ± SD, n = 8-12/group). Data from (A) and (C) are representative of three independent experiments, and data from (B) and (D) are pooled from three pigs. (**E**) Movat Pentachrome staining of mouse db/db wound sections on day 25 showing elastic fibers (black to blue/black), with dotted frames in overview images on the left magnified on the right. (**F**) Movat Pentachrome staining of diabetic pig wounds on day 16 showing elastic fibers (black to blue/black), with dotted frames in overview images on the left magnified on the right to depict the wound margin. Scale bars: 1 mm (left), 100 µm (right) in (A-F). \**P* < 0.05, \*\**P* < 0.01, \*\*\**P* < 0.001. K14, keratin 14.

**Fig. 4.**
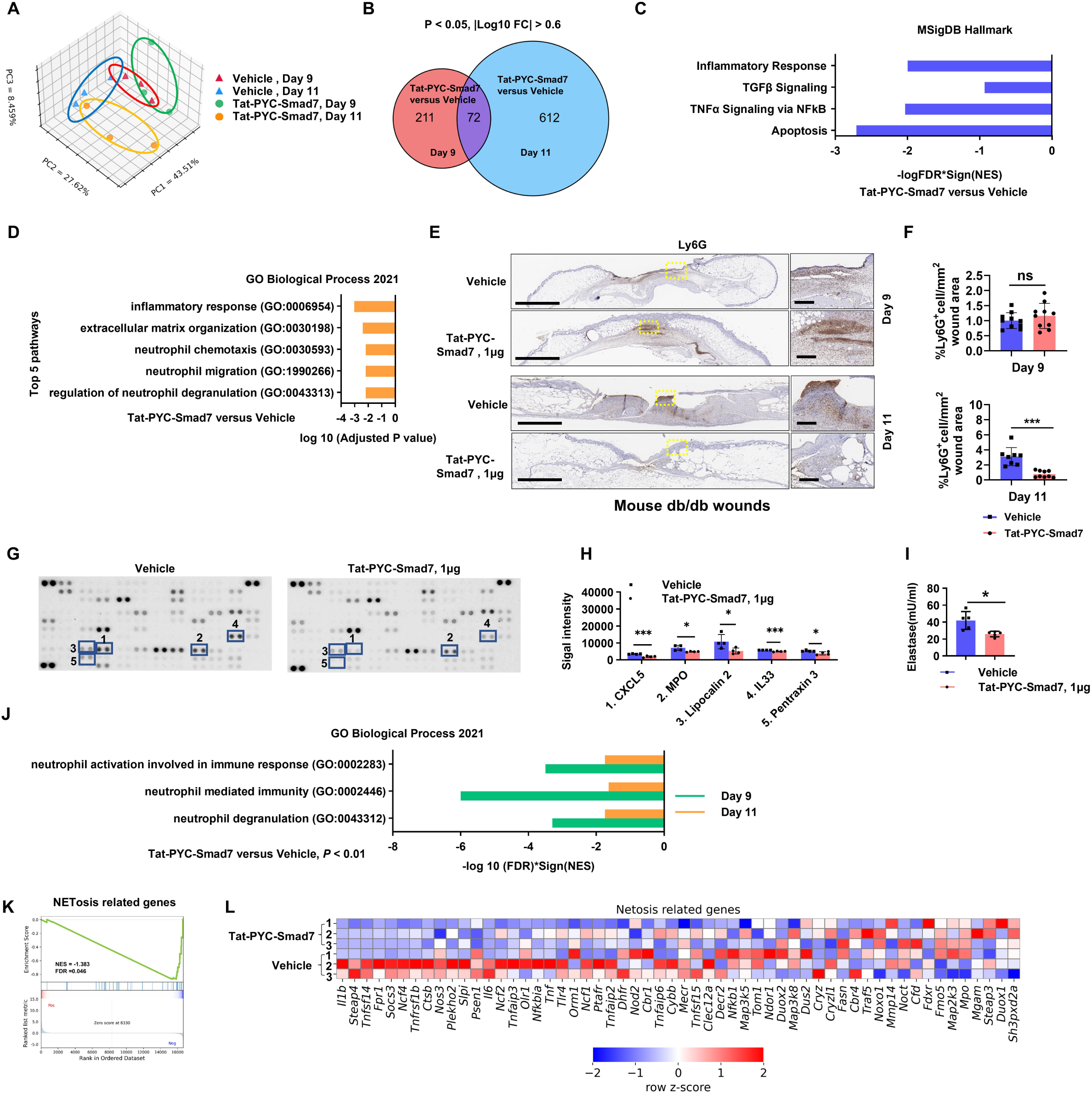
Transcriptional, histological and proteomic analysis showed that Tat-PYC-Smad7 had multiple targets and blunted neutrophil activities in diabetic wounds. (**A**) Principal component analysis of transcriptomic profiles from wound samples comparing Tat-PYC-Smad7 treatment versus Vehicle on days 9 and 11. (**B**) Venn diagram showing the overlap of DEGs between Tat-PYC-Smad7-versus Vehicle-treated wounds on day 9 and day 11. (**C**) GSEA of Hallmark gene sets for transcriptomes of wound samples from Tat-PYC-Smad7 versus Vehicle treatment on day 9 and day 11. (**D**) Top five enriched GO biological processes identified by Enrichr using DEGs from Tat-PYC-Smad7 versus Vehicle-treated wounds on days 9 and 11. (**E**) Representative IHC images of Ly6G^+^ neutrophils in wounds treated with Tat-PYC-Smad7 or Vehicle on days 9 and 11. Representative staining areas enclosed by dotted frames in overview images on the left are magnified in the right images. Scale bars: 1 mm (left), 100 µm (right). (**F**) Quantification of Ly6G^+^ neutrophils in wound tissues on day 9 and day 11 (mean ± SD, n = 8-10/group). (**G**) Representative proteomic array images of db/db wound lysates from mouse wounds treated with Tat-PYC-Smad7 versus Vehicle on day 11. (**H**) Quantification of signal intensity of selected inflammatory and neutrophil-associated proteins in (G) in Tat-PYC-Smad7-versus Vehicle-treated wounds (mean ± SD, n = 4/group). (**I**) Measurement of elastase activity in db/db wound lysates on day 11 (mean ± SD, n = 5/group). (**J**) GSEA of GO biological processes related to neutrophil activation, neutrophil-mediated immunity, and neutrophil degranulation in Tat-PYC-Smad7-treated wounds on days 9 and 11. (**K**) Enrichment plot and (**L**) heatmap of NETosis-related genes in Tat-PYC-Smad7-treated wounds compared to Vehicle-treated wounds on day 11. Data are representative of three independent experiments. ns indicates not significantly different, \**P* < 0.05, \*\**P* < 0.01, \*\*\**P* < 0.001. DEGs, differentially expressed genes; FC, fold change; GSEA, gene set enrichment analysis; GO, Gene Ontology; IHC, immunohistochemistry; MPO, myeloperoxidase; CitH3, citrullinated histone 3.

**Fig. 5.**
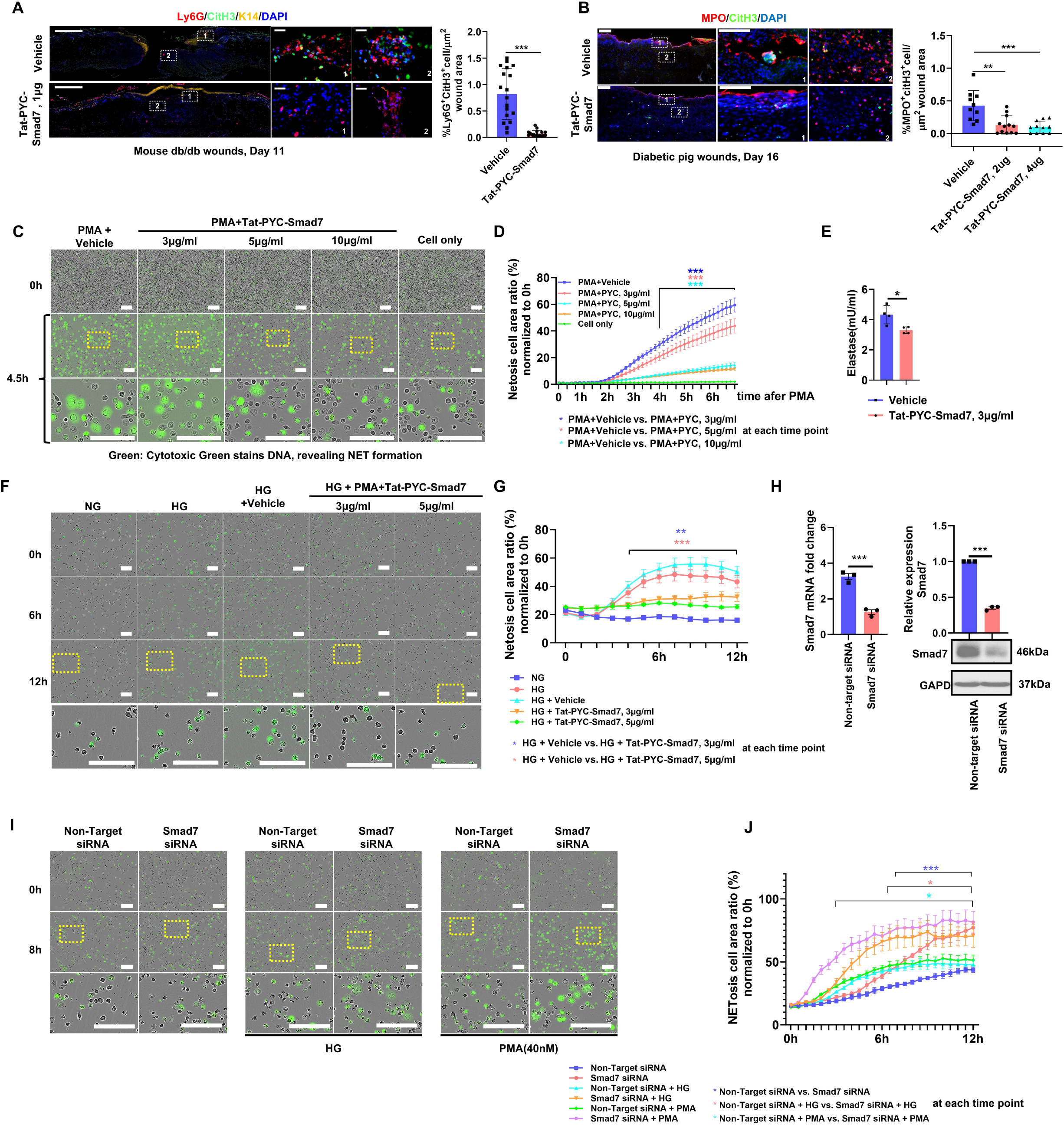
Tat-PYC-Smad7 blunted NETosis in dHL60 cells under both PMA- and HG-treated conditions. (**A**) Representative immunofluorescence images of mouse db/db wounds on day 11 stained for Ly6G, CitH3, K14, and DAPI with quantification of Ly6G^+^CitH3^+^ cells (mean ± SD, n = 17-18/group, pooled data from two independent experiment). (**B**) Representative immunofluorescence images of diabetic pig wounds on day 16 stained for MPO, CitH3, and DAPI with quantification of MPO^+^CitH3^+^ cells (mean ± SD, n = 10-12/group, pooled data from three individual pigs). (**C**) Real-time observation of NETosis using IncuCyte live-cell imaging in dHL60 cells treated with PMA and Tat-PYC-Smad7 or Vehicle at different concentrations, showing Cytotoxic Green fluorescence as a marker of extracellular DNA. (**D**) Quantification of NETosis from (C) based on Cytotoxic Green fluorescence (mean ± SEM, n = 4/group). (**E**) Elastase activity in PMA-induced dHL60 cells treated with Tat-PYC-Smad7 or Vehicle (mean ± SD, n = 4/group). (**F**) Time-lapse imaging of HG-induced NETosis in dHL60 cells treated with Tat-PYC-Smad7 or Vehicle. (**G**) Quantification of HG-induced NETosis from (F) (mean ± SEM, n = 4/group). (**H**) Smad7 knockdown efficiency confirmed by qPCR, Western blot, and quantification of band intensity (mean ± SD, n = 3/group). (**I**) Real-time observation of NETosis in Smad7-knockdown dHL60 cells under untreated, HG, or PMA stimulation. (**J**) Quantification of NETosis progression from (I) (mean ± SEM, n = 4/group). Representative staining areas enclosed by dotted frames are magnified in the corresponding images. Scale bars: 1 mm (left) in (A, B); 100 µm (right) in (B); 50 µm (right) in (A); 100 µm in (C, F, I); 20 µm (left), 10 µm (right) in (E). *In vitro* data are representative of three independent experiments. \**P* < 0.05, \*\**P* < 0.01, \*\*\**P* < 0.001. CitH3, citrullinated histone 3; MPO, myeloperoxidase; dHL60, differentiated HL-60; NG, normal glucose; HG, high glucose; siRNA, small interfering RNA.

### Tat-PYC-Smad7 promoted the healing of diabetic pig wounds

To assess the effect of Tat-PYC-Smad7 on wounds closely mimicking human skin, we used a miniature diabetic pig model ^19^. We used three diabetic pigs and punched twelve 2cm diameter full-thickness wounds in each pig for drug treatment (Fig.2A). Two doses of Tat-PYC-Smad7 protein (2 µg and 4 µg) or vehicle control of either hydroxypropyl cellulose (HPC) or hydroxyethyl cellulose (HEC) were topically applied every other day and wounds were covered with a medical dressing (Fig.2A). Immunohistochemistry staining for Smad7 revealed that Tat-PYC-Smad7 was detected in the epidermis and stromal cells at the wound edge in protein-treated wounds, while it was absent in vehicle-treated wounds (Sup Fig.6A). To assess whether topically applied Tat-PYC-Smad7 enters the bloodstream, a Smad7-specific ELISA was developed, capable of detecting levels as low as 5 ng/mL. Smad7 remained undetectable in systemic circulation in all treated pigs (Sup Fig.6B). We evaluated whether Tat-PYC-Smad7 induced an anti-drug antibody (ADA) response that could neutralize its therapeutic effects and found no detectable ADA against Tat-PYC-Smad7 in treated pigs (Sup. Fig 6C). Wound healing kinetics among individual pigs varied, resembling variations in human diabetic wounds ^31^, hence wound healing response to treatment of each pig by gross evaluation was analyzed individually. Grossly, the treatment effects of Tat-PYC-Smad7 (compared to vehicle) became evident from day 8, 5 and 11 for pig 1, 2 and 3, respectively (Fig.2B-C). Histological analysis indicated that more than 30% of the wounds were healed in the Tat-PYC-Smad7-treated group compared to 8% of the vehicle group (Fig.2D-E). Tat-PYC-Smad7 significantly reduces the remaining wound length as a percentage of the initial wound length compared to the vehicle control, indicating enhanced wound closure in diabetic pigs (Fig.2F). Moreover, Tat-PYC-Smad7 treatment resulted in longer keratinocyte migrating tongues in all three pigs compared to vehicle controls (Fig.2G).

**Fig. 6.**
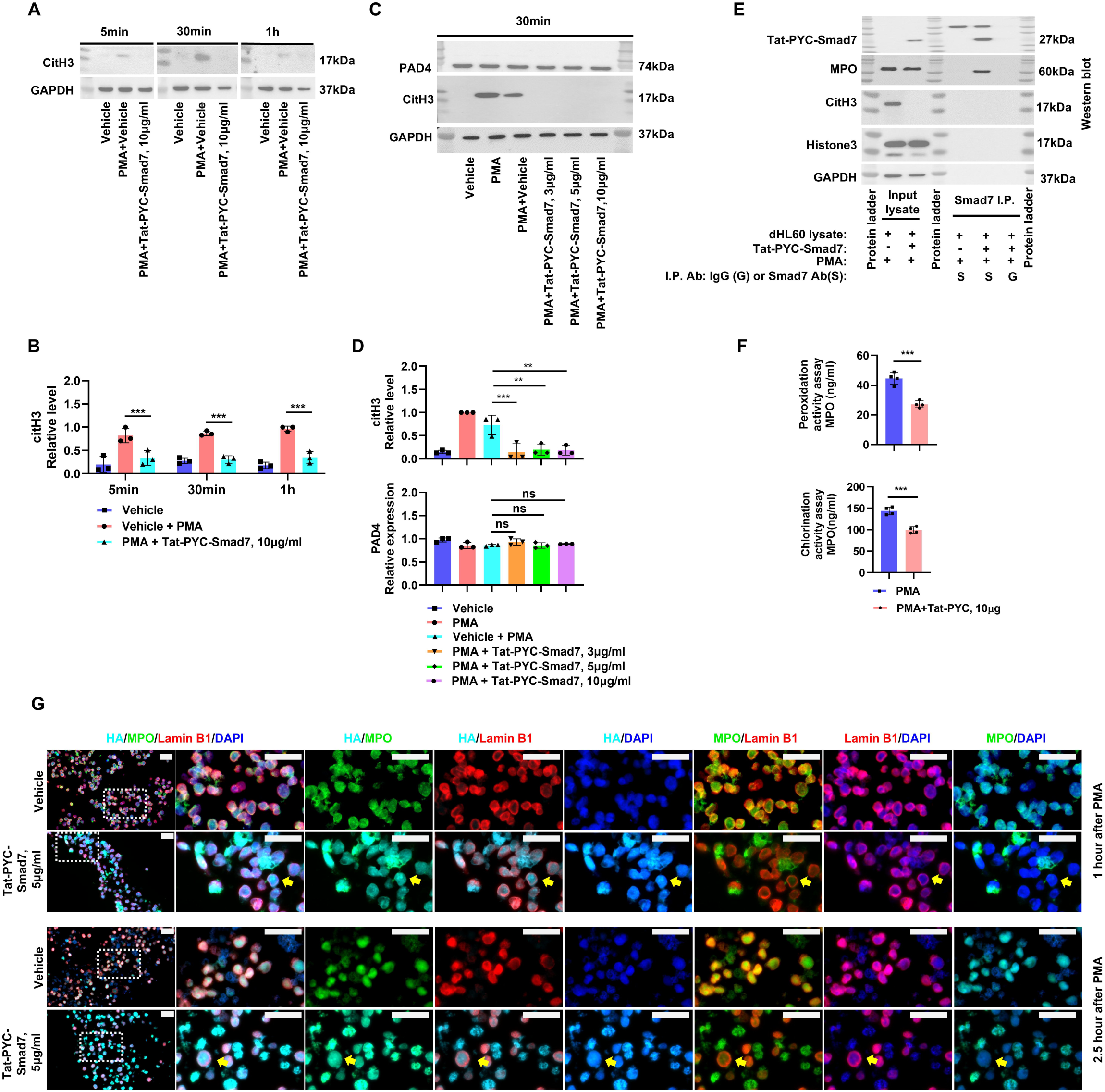
Tat-PYC-Smad7 bound MPO, reduced MPO nuclear entry and activity and suppressed NETosis. (**A**) Western blot analysis of CitH3 in dHL60 cells treated with PMA and Tat-PYC-Smad7 or Vehicle at different time points (5 min, 30 min, 1 h), with GAPDH as a loading control. (**B**) Quantification of CitH3 band intensity from three independent experiments (mean ± SD). (**C**) Western blot analysis of CitH3 and PAD4 in dHL60 cells treated with PMA and different concentrations of Tat-PYC-Smad7 (3, 5, 10 µg/mL), with GAPDH as a loading control. (**D**) Quantification of CitH3 and PAD4 band intensity from three independent experiments (mean ± SD). (**E**) Immunoprecipitation of cell lysates using an anti-Smad7 antibody or an isotype control antibody, followed by Western blot detection of associated proteins, including MPO, CitH3, and histone 3. (**F**) MPO activity assays measuring peroxidation and chlorination activities in dHL60 cells treated with Tat-PYC-Smad7 or Vehicle during PMA-induced NETosis (mean ± SD, n = 4 independent experiments). (**G**) Representative immunofluorescence images of dHL60 cells treated with Vehicle or Tat-PYC-Smad7 (5 μg/mL) for 10 minutes, followed by PMA stimulation (100 nM) for 1 hour or 2.5 hours. Cells were stained for MPO (green), HA-tagged Tat-PYC-Smad7 (cyan), Lamin B1 (red, nuclear membrane marker), and nuclei (DAPI, blue). The areas enclosed by the dotted boxes are magnified and shown on the right. Yellow arrows indicate cells in which MPO entered the nucleus, where colocalization of MPO with DNA and Tat-PYC-Smad7-HA was observed within a continuous Lamin B1 ring. Scale bars: 20 μm. ns, not significant; \*\**P* < 0.01; \*\*\**P* < 0.001. CitH3, citrullinated histone 3; MPO, myeloperoxidase; PAD4, peptidylarginine deiminase 4; PMA, Phorbol 12-myristate 13-acetate; dHL60, differentiated HL-60 cells; I.P., immunoprecipitation.

To understand the pathogenic targets of Tat-PYC-Smad7 in wound healing, we performed immunofluorescent staining of diabetic pig and db/db mouse wound sections and found that Tat-PYC-Smad7-treated wounds contain more Ki67^+^ cells and less αSMA^+^ area compared to vehicle-treated wounds (Fig.3A-B, Sup Fig.7, suggesting increased keratinocyte proliferation and less fibrotic response. Moreover, Tat-PYC-Smad7 restored angiogenesis in the stroma area under the migrated epidermis in diabetic pig wounds and db/db mouse wounds, with more CD31-positive areas at the edge of Tat-PYC-Smad7-treated wounds compared to control wounds (Fig.3C-D, Sup Fig.7B). Additionally, Tat-PYC-Smad7-treated wounds had more regenerative and organized wavy elastic fibers (stained as black to blue/black striae) entwined among collagenous fibers shown by Pentachrome staining in healed wounds of db/db mice on day 25 and diabetic pig wounds on day 16 (Fig.3E-F), indicating that Tat-PYC-Smad7 improved ECM deposition and collagen remodeling.

To understand the molecular basis of Tat-PYC-Smad7 effects, we performed RNA-seq on db/db mouse wound samples excised at identical locations treated with vehicle or Tat-PYC-Smad7 on days 9 and 11. Days 9 and 11 are critical stages at which db/db wounds start to transition towards a chronic state, compared to normal wounds ^18^. Principal components analysis revealed distinct transcriptomic profiles of Tat-PYC-Smad7-treated wounds vs vehicle-treated wounds (Fig.4A). The number of differentially expressed genes (DEGs) was 211 on day 9 and 612 on day 11, with 72 DEGs overlapping between the two timepoints (Fig.4B). Gene set enrichment analysis (GSEA) using the Hallmark gene sets showed that Tat-PYC-Smad7 inhibited TGFβ and NFκB signaling, inflammatory responses and apoptosis pathways (Fig.4C and Sup Fig.8), consistent with known TGFβ/NFkB effects on wound healing in our previous reports ^21,32^ as well as phenotypic changes seen in Fig.3A-B. Interestingly, enrichment analysis using the overlapping DEGs between day 9 and day 11 on gene ontology (GO) terms identified that neutrophil functions affected by Tat-PYC-Smad7 were among three out of top five enriched biological processes (Fig.4D). As DEG enrichment analysis indicated a significant impact of Tat-PYC-Smad7 on neutrophils on both day 9 and day 11, we examined Ly6G^+^ neutrophils in the wound beds and found that the number of Ly6G^+^ cells was unchanged by Tat-PYC-Smad7 treatment on day 9, but significantly reduced on day 11 (Fig.4E-F), suggesting that Tat-PYC-Smad7 may affect neutrophil functionality and chemotaxis. Analysis of wound tissue lysates from day 11 using an antibody array revealed that wounds from mice treated with Tat-PYC-Smad7 had reduced proinflammatory proteins and cytokines, including CXCL5 ^33^, lipocalin2 ^34^, and cytokine IL33 ^35^, and granule components myeloperoxidase (MPO) ^36^ and Pentraxin 3 ^37^, compared to wounds from mice treated with vehicle (Fig.4G-H). Tat-PYC-Smad7-treated wounds also had decreased elastase production on day 11(Fig.4I), which is often associated with neutrophil degranulation or NETosis ^38^. Therefore, Tat-PYC-Smad7 exerts effects on both epidermis and stroma cells in diabetic wounds with better healing outcomes.

### Tat-PYC-Smad7 blunted neutrophil activities in diabetic wounds at molecular and cellular levels

In addition to the impact of Tat-PYC-Smad7 on keratinocytes and ECM regulation, GSEA of GO terms on day 9 and day 11 showed the top 3 functional clusters associated with neutrophil activities that were consistently attenuated by Tat-PYC-Smad7 treatment (Fig.4J, Sup Fig.9A). This is consistent with the later-stage neutrophil infiltration observed in db/db mice, as Ly6G staining showed only sparse neutrophil presence on days 3 and 5 in comparison to abundant neutrophil infiltration on day 9 and day 11 after wounding (Sup Fig. 9B). Immunohistochemistry staining of the wound tissues showed penetration of Tat-PYC-Smad7 in both keratinocytes and Ly6G^+^ cells in protein-treated wounds (Sup Fig.10), indicating a possible direct effect of Tat-PYC-Smad7 on neutrophil functions. Additionally, in diabetes, high glucose (HG) levels induce neutrophils to undergo NETosis ^15^. Neutrophils recruited to wound sites almost immediately undergo NETosis and complete NET release in ∼2.5 hours ^15^. To evaluate the Tat-PYC-Smad7 effect on NETosis *in vivo*, we examined a transcriptomic profile of NETosis ^39^ previously demonstrated to be specifically involved in diabetic wound healing defects ^15^. We found that the NETosis-related gene set was downregulated in wounds treated with Tat-PYC-Smad7 (Fig.4K and L), with similarly strong enrichment as other neutrophil-associated gene sets (Fig.4J and Sup Fig.9A). Furthermore, in both diabetic pig and db/db mouse models, immunofluorescence staining indicated fewer neutrophils undergoing NETosis in the Tat-PYC-Smad7-treated wound beds, determined by co-staining of Ly6G or MPO (neutrophil markers) and citrullinated histone 3 (citH3) (NETosis marker) with DAPI (DNA marker) (Fig.5A-B). These data suggest that Tat-PYC-Smad7 blunts neutrophil NETosis in diabetic wounds.

### Tat-PYC-Smad7 prevented PMA and high glucose-induced NETosis in dHL60 cells

To test if Tat-PYC-Smad7 directly affects NETosis, we performed an *in vitro* assay using phorbol myristate acetate (PMA)-induced NETosis in dimethyl sulfoxide (DMSO)-differentiated HL60 cells (dHL60), a neutrophil cell culture model ^40^. Tat-PYC-Smad7-HA was detected in the nucleus 5min after treatment, showing highly efficient cell penetration (Sup Fig.11A). We observed that dHL60 cells treated with Tat-PYC-Smad7 had fewer cells marked with NETs 4 hours post-induction with PMA (Sup Fig.11B). Live cell imaging using IncuCyte showed significant attenuation of PMA-induced NETosis by Tat-PYC-Smad7 in a dose and time-dependent manner (Fig.5C-D). Cytotoxic Green, a cell-impermeant DNA dye that labels extracellular DNA released during loss of membrane integrity, colocalized with citH3-positive neutrophils and MPO in PMA-induced dHL60 cells (Sup Fig. 12), confirming that the observed signal represented NETosis. We also observed reduced elastase release from dHL60 cells 4 hours after PMA induction following Tat-PYC-Smad7 treatment, which is considered an additional marker of NETosis (Fig.5E). NETosis induced by HG, mimicking the blood glucose levels in diabetic wounds, was also inhibited by Tat-PYC-Smad7, (Fig.5F-G). Furthermore, siRNA knockdown of endogenous Smad7 in dHL60 cells significantly increased NETosis under both PMA- and HG-induced conditions (Fig.5H-J), suggesting Smad7 suppresses NETosis. PMA-induced NETosis was not affected by BSA, TGFβ or TGFβ inhibitor treatments (Sup Fig.13), suggesting attenuation of NETosis by Tat-PYC-Smad7 was not related to its effect of antagonizing TGFβ signaling or as a non-specific protein.

We next explored the molecular mechanism underlying the effect of Tat-PYC-Smad7 on neutrophil activity. As histone citrullination is the critical step for NETosis ^41^, we first investigated whether Tat-PYC-Smad7 affects this process. PMA-induced citrullinated histone 3 (citH3) was blocked by Tat-PYC-Smad7 treatment at all time points examined (Fig.6A-B) and all doses of Tat-PYC-Smad7 blocked citrullination of histone 3 (Fig.6C-D). PAD4, the key enzyme for histone citrullination ^41^, showed no significant change in protein levels following Tat-PYC-Smad7 treatment (Fig.6C-D). We conducted a pulldown assay with a Smad7 antibody recognizing Tat-PYC-Smad7 using lysates of PMA-treated dHL60 cells treated with the vehicle or Tat-PYC-Smad7. Mass spectrometry analysis showed several proteins were enriched in Tat-PYC-Smad7 samples pulled down with anti-Smad7 antibody (Sup Fig.14A-B). A list of the top Tat-PYC-Smad7-binding targets includes MPO (Sup Fig.14B), a protein required for NETosis via translocation into the nucleus to promote chromatin unfolding ^41, 42^. Consistent with mass spectrometry findings, MPO was detectable in anti-Smad7 pulldown samples only from Tat-PYC-Smad7/PMA-treated dHL60 cell lysates, supporting a potential interaction between Tat-PYC-Smad7 and MPO (Fig.6E). Functional assays revealed that Tat-PYC-Smad7 treatment attenuated both peroxidation and chlorination activities of MPO under NETosis condition (Fig.6F), which contribute to NETosis through reactive oxidant generation, lipid peroxidation, and chromatin remodeling ^43^. To further investigate the impact of Tat-PYC-Smad7 on MPO dynamics during NETosis, we performed immunofluorescence co-staining with MPO, HA-tagged Tat-PYC-Smad7 and Lamin B1 (a nuclear membrane marker). MPO enters the nucleus 1 hour post-PMA treatment in the vehicle condition, but MPO is largely excluded from the nucleus in the Tat-PYC-Smad7-treated cells at this timepoint (Fig. 6G). By 2.5 hours, Tat-PYC-Smad7-treated cells still showed reduced nuclear MPO accumulation and extracellular spread of MPO compared with vehicle control (Fig. 6G). We also observed instances where MPO entered the nucleus of treated cells, where it colocalized with DNA (DAPI) and Tat-PYC-Smad7-HA within a continuous Lamin B1 ring at 1 hour and 2.5 hours of PMA-induced NETosis (Fig. 6G). Consistently, at 2.5 hours, colocalization between Tat-PYC-Smad7 with MPO appeared mainly in MPO-positive nuclei in Tat-PYC-Smad7-treated and PMA-stimulated dHL60 cells (Sup Fig.15). Colocalization analysis showed that about 20-30% of the strong HA and MPO signals appeared in the same locations, based on thresholded Manders’ coefficients. The thresholded Pearson’s correlation coefficient (Rcoloc) was close to +1, indicating that HA and MPO signals were strongly associated in signal-rich regions (Sup table 2).

## Discussion

### Smad7 has a direct effect on promoting healing at the diabetic wound site

In this study, we showed that Smad7 promoted healing of diabetic wounds in mice and pigs. K5.Smad7 transgene expression ameliorated diabetic phenotypes in db/db mice, including improvements in body weight and blood glucose levels (Fig.1B-C). As K5.Smad7 mice overexpress Smad7 in stratified epithelia ^24^, this protective effect may derive from local homeostatic alterations, potentially implicating metabolic processes ^44^. However, Tat-PYC-Smad7 topical treatment to diabetic wounds promoted healing without reducing body weight and glucose levels, and the improvement to wound healing in diabetic pigs was compared to vehicle controls in the same pig (same glucose and body weight). Taken together, Smad7 effect on wound healing is primarily due to its direct and local effects.

Our previous work demonstrated TGFβ/NFκB signaling activation in human diabetic wounds and showed that full-length Smad7-based protein could promote wound healing defects in K5.TGFβ mice, a model that primarily reflects inflammatory skin pathology rather than diabetes ^45^. In the present study, we directly studied clinically relevant diabetic wound models in both mouse and pig models, and assessed therapeutic efficacy by Tat-PYC-Smad7.Although only containing partial Smad7, PYC-Smad7 proved to have critical functional domains of Smad7 for promoting wound healing. Moreover, our data reveal a previously unrecognized effect of Smad7 on NETosis. These findings highlight that Smad7 promotes repair primarily through local mechanisms and further support the translational potential of Tat-PYC-Smad7 as a therapeutic strategy.

### Smad7-based treatment is a novel approach for diabetic wound healing that elicits multiple local therapeutic effects compared to Regranex

In comparison to Regranex, a growth factor that predominantly stimulates fibroblasts to increase granulation formation and promote wound closure, Smad7-based biologics accelerated reepithelization because it reduced TGFβ-induced growth inhibition and cell cycle arrest in keratinocytes, and promoted keratinocyte migration ^22, 45, 47^. In addition, Smad7 is not a secreted protein, and its local, short-term delivery did not exert systemic effects, which is likely due to its rapid entry into local cells and low doses needed for efficacy ^48^. In contrast, Regranex, a mitogen, strongly boosts myeloid cell proliferation and activation, recruiting immune cells for tissue repair ^13^. However, overstimulating the immune system could exacerbate chronic inflammation that greatly hinders diabetic wound healing ^49^. To support this notion, several cytokines elevated by Regranex - including Angiopoietin-2, CXCL16, CXCL5, and LDL R - are linked to chronic inflammation and fibrosis conditions^28, 29, 30^. Elevated LDL R, associated with atherosclerosis, aligns with the foam-like cells observed in Regranex-treated wounds, resembling those in atherosclerotic plaques and suggesting active inflammation with profibrotic characteristics^27^. In contrast, because TGFβ is a central mediator of fibrosis ^50^, the anti-TGFβ function of Tat-PYC-Smad7 confers an anti-fibrotic effect, leading to better healing outcomes with less scarring and more tissue regeneration in both mouse and pig models. This is evidenced by the reduced αSMA^+^ area in db/db mouse wounds treated with Tat-PYC-Smad7 (Fig.3A). In the diabetic pig wound model, all three pigs responded to the Tat-PYC-Smad7 treatment. The treatment responding rate exceeded the 28% efficacy rate observed with Regranex in clinical studies ^5^. Furthermore, our previous studies show that topical application of a Smad7-based drug did not increase the risk of carcinogenesis that can occur with growth factor therapies ^46, 51^ and could be better tolerated during repeated dosing and long-term usage. Therefore, Tat-PYC-Smad7 may have better treatment efficacy and potential benefits over Regranex as a therapeutic option for diabetic wounds.

### Tat-PYC-Smad7 targets wound epidermis and stroma to promote the healing of diabetic wounds and better healing outcomes

In this study, Tat-PYC-Smad7 showed multiple functional impacts on keratinocytes, ECM and inflammation in diabetic wound models. Inhibition of TGFβ/NFκB signaling could explain the effect of Tat-PYC-Smad7 on promoting keratinocyte survival and ECM organization, consistent with our previous studies ^21, 32, 45, 46, 47^. Further, our *in vitro* studies suggested that Tat-PYC-Smad7 could target neutrophils by blocking NETosis. Although Tat-PYC-Smad7 suppressed MPO enzymatic activity, this may not fully account for its inhibitory effect on NET formation, as MPO peroxidase activity may not be essential for NETosis^43^. Nevertheless, MPO is known to facilitate chromatin decondensation during NETosis independent of its enzymatic function ^52, 53^, and its nuclear presence is required for efficient NET formation ^43, 52^. Our imaging data showed that Tat-PYC-Smad7-treated cells exhibited reduced MPO nuclear accumulation and extracellular spread compared with vehicle control, suggesting that Tat-PYC-Smad7 may prevent MPO nuclear translocation. In addition, in a subset of treated cells where MPO did enter the nucleus, MPO is colocalized with DNA and Tat-PYC-Smad7-HA within an apparently intact nuclear membrane, indicating that such interactions can occur prior to overt nuclear envelope disruption. Together, these results suggest that Tat-PYC-Smad7 may exert stepwise effects in disrupting NETosis via reducing MPO nuclear translocation and chromatin decondensation. However, the mechanisms regulating MPO nuclear translocation and chromatin decondensation during NETosis remain incompletely understood. Elucidating this process and the precise role of Tat-PYC-Smad7 in regulating the mechanism of NETosis, warrant future studies. Importantly, as NETosis is a major source of proinflammatory cytokines, proteases, and reactive oxygen species (ROS) in chronic wounds^12^, inhibition of NETosis by Tat-PYC-Smad7 likely contributes to reduced inflammation independent of changes in neutrophil infiltration. This is consistent with our in vivo findings, where Tat-PYC-Smad7 treatment reduced inflammatory cytokine expression and ROS levels prior to a significant decrease in neutrophil numbers (Fig.4E-F).

Previous studies have highlighted the detrimental impact of prolonged neutrophil activation in the disruption of re-epithelialization, angiogenesis, and ECM destruction in chronic wounds^7^. The accelerated re-epithelialization in Tat-PYC-Smad7-treated diabetic wounds, angiogenesis, and ECM remodeling could be partially attributed to neutrophil inhibition by Tat-PYC-Smad7. In addition to a direct effect, Tat-PYC-Smad7 may affect epidermal signals/stimuli that activate neutrophils at the wound site to further curb neutrophil activities. For example, Tat-PYC-Smad7 may regulate the neutrophil responses to Toll-like receptors or NOD-like receptors by downregulating keratinocyte-derived pathogen-associated molecular patterns or damage-associated molecular patterns via NFκB signaling inhibition ^54^. Therefore, the effects of Tat-PYC-Smad7 on both the epidermis and stroma may synergistically enhance wound healing, leading to improved treatment outcomes.

Our study indicates that Tat-PYC-Smad7 affects multiple signaling cascades involving TGFβ/NFκB and NETosis in wound healing defects, making it a potent multi-functional protein agent. Regulating NETosis by topical Tat-PYC-Smad7 application could minimize off-target side effects compared to systemic NET inhibition^15, 55^. Future studies will identify signaling pathways and molecular mechanisms leading to the binding of Smad7-based biologics with MPO and downstream mechanistic steps of NETosis. In addition, as wound healing is a complex process requiring the coordination of multiple cell types and events, the pleiotropic effects of Tat-PYC-Smad7 targeting both local epithelial and proinflammatory cells could offer potential advantages over therapeutics targeting a single event or specific pathway. While our findings indicate that Tat-PYC-Smad7 selectively modulates neutrophil activity without broadly impairing neutrophil recruitment, future investigations are necessary to assess its potential impact on host defense mechanisms in infected wounds. In conclusion, our results demonstrate the potential of a Smad7-based biologic to treat chronic diabetic wounds. The mechanisms of action include inhibition of proinflammatory pathways in the epidermis, promotion of keratinocyte migration for rapid reepithelization and ECM remodeling, and blunted NET formation and MPO activity.

## Methods

Detailed methods are included in the supplementary methods. The primary antibodies used are detailed in Sup. Table 3.

### K5.Smad7/db/db bigenic mice

The Institutional Animal Care and Use Committee at the University of Colorado Denver Anschutz Medical Campus and the University of California Davis Medical Center approved all described animal experiments. K5.Smad7/db/db bigenic mice were generated by breeding previously established K5.Smad7 mice ^24^ with heterozygous B6.BKS(D)-Leprdb/J mice (JAX000697) or heterozygous BKS.Cg-Dock7m+/+Leprdb/J mice (JAX000642). The genotyping of the bigenic mice was performed as previously described^24, 56^. No gender differences were observed in our previous wound study ^21^. The female mice were utilized once they reached 8-10 weeks of age. Before the wound healing test, mice were weighed, and their blood glucose levels were assessed.

### Tat-PYC-Smad7 treatment preparation

Tat-PYC-Smad7 was generated and formulated as previously described^48^. Tat-PYC-Smad7 is a recombinant protein that contains the PY and C domains of Smad7 (203-426aa) with a cell-permeable Tat tag ^21, 51^. An optional HA epitope at the C-terminus of Tat-PYC-Smad7 enabled tracking of the protein. HPC (F42, 1% Klucel HF, Ashland) containing PBS and 30% glycerol or HEC (1% Natrosol 250HX, Ashland) containing PBS and 30% glycerol gel was used as the base gel. Vehicle and Tat-PYC-Smad7 gel were formulated by carefully mixing protein buffer or protein stock with the base gel at a 1:1 ratio. All doses of the formulation were aliquoted and stored at -80°C. Prior to the start of dosing, dose formulations were freshly thawed and gently inverted to mix.

### db/db mouse wound model

8-12 week-old female BKS.Cg-Dock7m+/+Leprdb/J (db/db) mice (JAX000642) were purchased from Jackson Laboratory. Mice were anesthetized with 5% inhaled isoflurane, and the dorsal surface was shaved. Two pairs of full-thickness dorsal wounds (6 mm) were created to remove epidermis, dermis, and panniculus carnosus. Tat-PYC-Smad7 or vehicle gel (0, 0.5, 1 μg) was applied (10 μL per wound), while Regranex was applied using a cotton swab. Wounds were covered with Tegaderm film. Treatments were given every other day from day 1 to endpoint (days 3, 7, 8, 9, 11, or 25). Wound areas were calculated using Python-based image analysis. Wound tissues were collected for histological evaluation, lysis, or RNA extraction.

### Diabetic porcine wound model and Tat-PYC-Smad7 treatment

Diabetic porcine wound model was established by Sinclair Research Center, LLC (currently Altasciences-Columbia pre-*Clinical*) following their approved animal protocols (Auxvasse, MO). Yucatan mini-pigs (castrated male 27-30kg) were used for the experiments. Diabetes was induced using an established method ^42^ to achieve fasting blood glucose levels of 150mg/dL or above at study enrollment. On day 1, animals were anesthetized and intubated with 5% isoflurane. Twelve 2 cm full-thickness excisional wounds (6 per side) were created along the dorsal spine using a skin punch. Hemostasis was achieved with epinephrine-soaked gauze (1:10,000 in sterile saline). After cleaning, each wound was covered with Tegaderm film and administered Tat-PYC-Smad7 or vehicle gel through the film, followed by a second Tegaderm layer and adhesive dressing. Foam pad and vet wrap secured the dressings. Dosing occurred every other day, with adjustments from 0.5 mL to 0.1 mL per wound. Wounds were photographed and analyzed, and animals were sacrificed two hours post-final dose. Wound tissue and serum samples were collected for further histological evaluation or assays.

### Cell culture, treatment and siRNA transfection

HL60 cells (University of Colorado Cell Technologies Shared Resource) were cultured in RPMI 1640 medium supplemented with 10% FBS and 1% Primocin™. dHL60 cells were differentiated with culture media supplemented with 1.3% DMSO (Sigma, D2650) as previously described ^40^. For PMA-induced NETosis experiments, cells were pretreated for 10 min with size exclusion chromatography (SEC) buffer (250 mM NaCl, 25 mM NaPO_4_, pH 7.6) (vehicle control), Tat-PYC-Smad7 or BSA (3, 5, or 10 µg/mL in SEC buffer) or incubated for 48 hr with TGFβ1 (2 or 5 ng/mL) or TGFβ inhibitor LY2109761 (5 ng/mL) prior to PMA (20 or 100 nM) stimulation. For high-glucose studies, dHL60 cells were cultured in F12 medium containing normal glucose (7 mM) with 26 mM mannitol as osmotic control ^15^ or HG (33 mM, for 12 hr, with or without Tat-PYC-Smad7 (3 or 5 µg/mL). For siRNA knockdown, dHL60 cells (300,000 cells/well) were transfected with 0.02 nmol Smad7 siRNA or non-targeting control using 1.2 µL Lipofectamine™ RNAiMAX. After 48 hr, cells were harvested for RT-PCR/Western blot analysis or subjected to NETosis induction (HG or PMA) with subsequent IncuCyte monitoring.

## Abbreviations

ECM: extracellular matrix
PDGF: platelet-derived growth factor
ROS: reactive oxygen species
HPC: hydroxypropyl cellulose
HEC: hydroxyethyl cellulose
DEGs: differentially expressed genes
GSEA: Gene set enrichment analysis
GO: gene ontology
MPO: myeloperoxidase
PMA: phorbol myristate acetate
DMSO: dimethyl sulfoxide
dHL60: differentiated HL60 cells
BSA: bovine serum albumin
citH3: citrullinated histone 3
SEC: size exclusion chromatography

## Data Availability

RNA-seq data are available in GEO (GSE274513). Mass spectrometry data are in MassIVE (MSV000095545; temporary access: username "MSV000095545_reviewer", password "a"). These credentials are provided for reviewer access, and the dataset is currently private. For any issues accessing the data, please contact the corresponding author.

## Acknowledgments

This work was funded by National Institutes of Health grants DE024659, DE028718, and AR078669 to XJW and CDY. YK received funding through the Milstein Medical Asian American Partnership Foundation Awards in Dermatology and a Sequencing Plot Award from the Genomics and Microarray Core at the University of Colorado Denver. We thank Dr. David Orlicky for pathology support. We thank Sinclair Research Center and Sinclair BioResources for the establishment of diabetic pig models and treatment tests. We thank Ivan Lu in the Immunology Graduate Group at UC Davis for his assistance with HL60 cell culture and Yi Wang for her assistance with organizing the gross images of the wounds We also extend our appreciation to Qian Chen from the Center for Genomic Pathology Laboratory (CGPL) at UC Davis for her timely help with cell pellet embedding. Additionally, we thank Dr. Ingrid Brust-Mascher from the Advanced Imaging Facility at the UC Davis School of Veterinary Medicine for her expert support with confocal imaging. We extend our appreciation to several entities for their valuable contributions: the University of Colorado Cancer Center Cell Technologies Shared Resource for their assistance with cell culture and use of the IncuCyte, the Mass Spectrometry Proteomics Shared Resource Facility at the University of Colorado for their mass spectrometry proteomic analyses, the Gates Center for Regenerative Medicine at the University of Colorado’s Anschutz Medical Campus for their histological support, and the RNA Biosciences Initiative at the University of Colorado for their support with RNA sequencing. Additionally, we acknowledge the Biostatistics and Bioinformatics Shared Resource at the University of Colorado Cancer Center for their assistance with processing RNAseq data. These shared resources are supported by the University of Colorado Cancer Center Support Grant (P30CA046934).

## Author contributions

YK, CDY, XJW designed experiments. YK performed the experiments, analyzed the data, and wrote the paper. BZL performed the bioinformatics analysis. FL performed some experiments and analyzed the data. RKR performed ELISA and ADA assays and analyzed data. DW and SW purified Tat-PYC-Smad7 protein. SW supported animal experiments. STH, SS and SRC assisted with editing. CDY and XJW also supervised the entire project and participated in writing and editing.

