## Supplementary tables for "Smad7-based biologic targeting epidermis and stroma promotes healing of diabetic wounds in mice and pigs"

### Supplementary Materials

### Sup Table 1. Demographics of the mice across genotypes in Fig.1B and C.

| Genotype | WT | | db/WT | | db/db | | K5.Smad7 | | K5.Smad7/db/db | |
|---|---|---|---|---|---|---|---|---|---|---|
| Sex | Female | Male | Female | Male | Female | Male | Female | Male | Female | Male |
| Number of mice | 3 | 9 | 4 | 10 | 11 | 5 | 5** | 1 | 4 | 6 |
| Mean Body weight (g)±SD | 19±1.1 | 25±5.3* | 22±2.3 | 28±2.5* | 51±5.7 | 52±5.7 | 18±2.2 | 27 | 35±4.1 | 41±7.2 |
| Mean Glucose (mg/dL)±SD | 182±35 | 151±41 | 136±24 | 175±33 | 456±103 | 416±111 | 126±16 | 146 | 270±144 | 264±90 |

*: $P < 0.05$ compared to male and female mice; **: Glucose was measured in 3 females

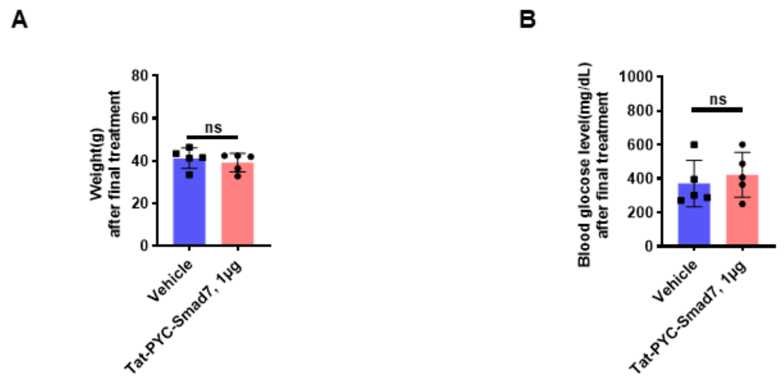


**Sup Fig.1. Topical application of Tat-PYC-Smad7 did not significantly affect blood glucose and body weight compared to the Vehicle.** Body weight **(A)** and blood glucose levels **(B)** of db/db mice after the final treatment of Tat-PYC-Smad7/Vehicle in the wounds on day 11. 5 samples from each group were analyzed using unpaired t test. Data are representative of independent wound experiments with 5 mice per group. One data point represents one mouse. Data represents the mean ± SD. ns indicates not significantly different.

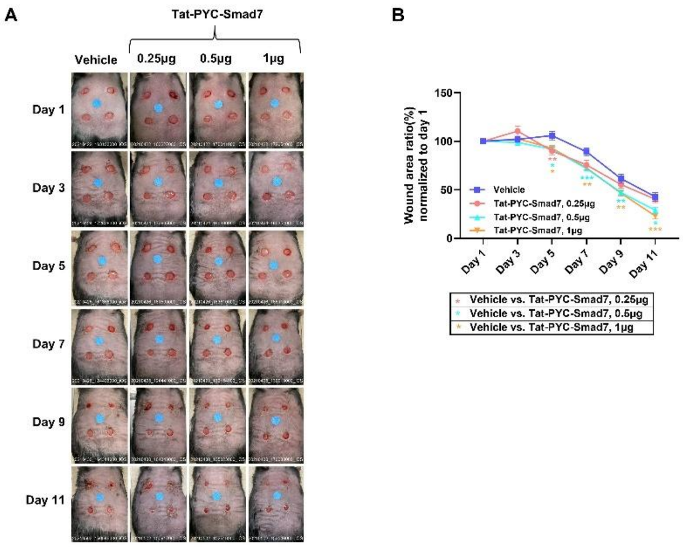


**Sup Fig.2. Tat-PYC-Smad7 promoted the db/db wound healing in a dose-dependent manner.** Representative gross images (Colored dot stickers with 6 mm in diameter were placed at the center of each wound served as a scale reference). **(A)** and quantification of the wound area **(B)** of db/db wounds treated with different dose of the Tat-PYC-Smad7 or Vehicle treatment. 20-24 wound samples from each group were quantified using two-way ANOVA with Tukey's multiple comparison testing. Data are representative of at least 3 independent experiments with 4 to 5 mice per group in each. Data represents the mean ± SEM. *$P$ <0.05, **$P$ < 0.01, ***$P$ < 0.001.

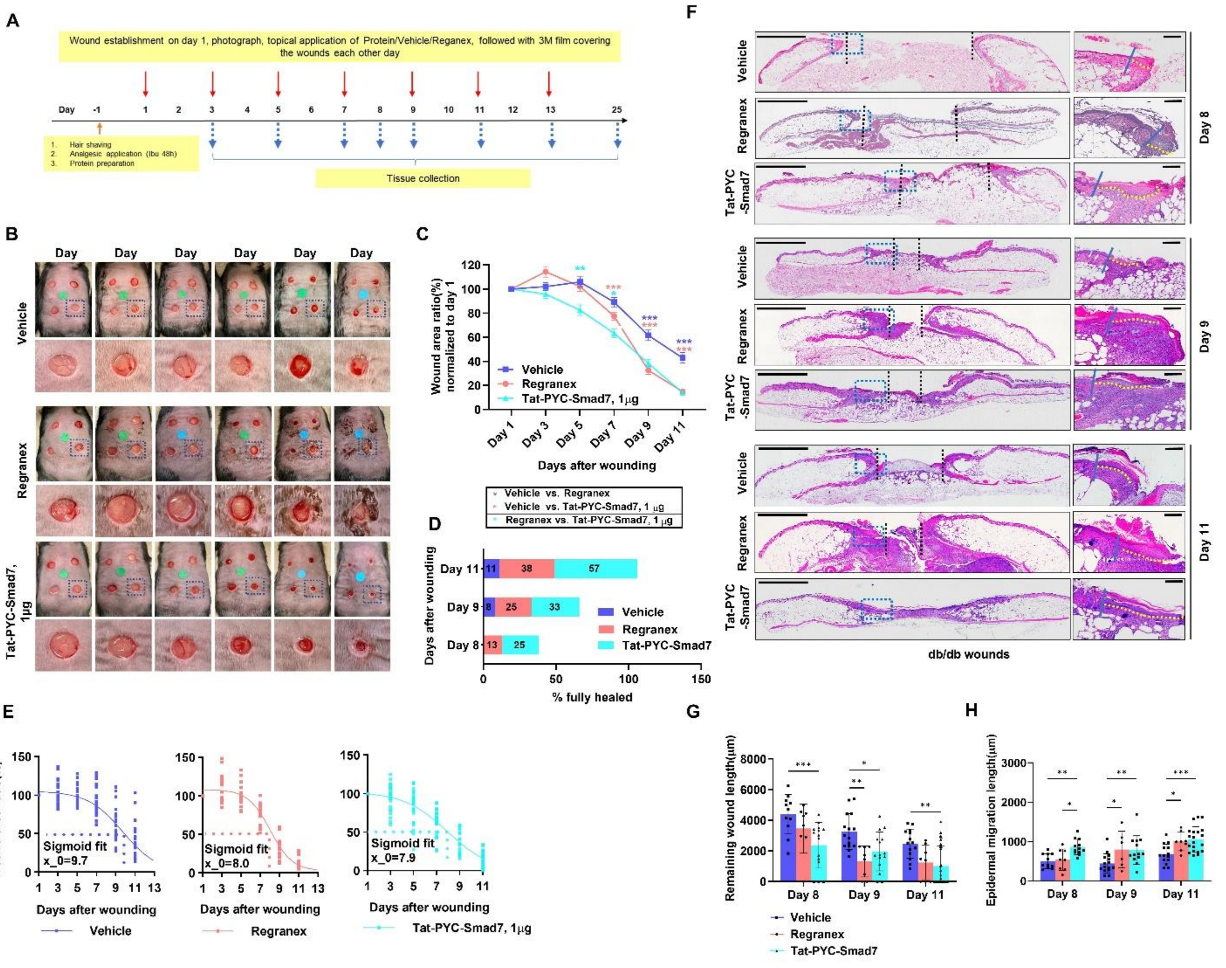


**Sup Fig.3. Topical application of Tat-PYC-Smad7 promoted diabetic wound healing better than Regranex.** **(A)** Experimental design for wounding and Vehicle/Tat-PYC-Smad7/Regranex treatment in female db/db mice. Representative gross images (Colored dot stickers with 6 mm in diameter were placed at the center of each wound served as a scale reference) **(B)**, quantification of the wound area **(C)**, and wound healing rate **(D)** of db/db wounds. **(E)** Sigmoid curve fit showing real-time gross quantification of the wound area. **(F)** Representative images of H&E-stained skin sections with quantification of remaining wound length **(G)** and epidermal migration length **(H)** at the indicated time points after wounding. The "dermabond-like" appearance in Regranex-treated groups in panel B is the residual marks left by the application of Regranex, which is an adhesive gel with high viscosity. Areas enclosed by the dotted dark blue boxes are magnified and shown below or to the right. Dotted yellow lines indicate migrating tongue. Dotted black lines indicate the edge of the wounds. Scale bars: 2mm (left), 200μm (right) in (F); 16-20 wound samples from each group were quantified using two-way ANOVA with Tukey's multiple comparison testing (C, G and H). Data are representative of at least 3 independent experiments with 4 to 5 mice per group in each. Data represents the mean ± SEM in (C). the mean ± SD in (G and H). **P* <0.05, ***P* < 0.01, ****P* < 0.001. H&E, hematoxylin and eosin.

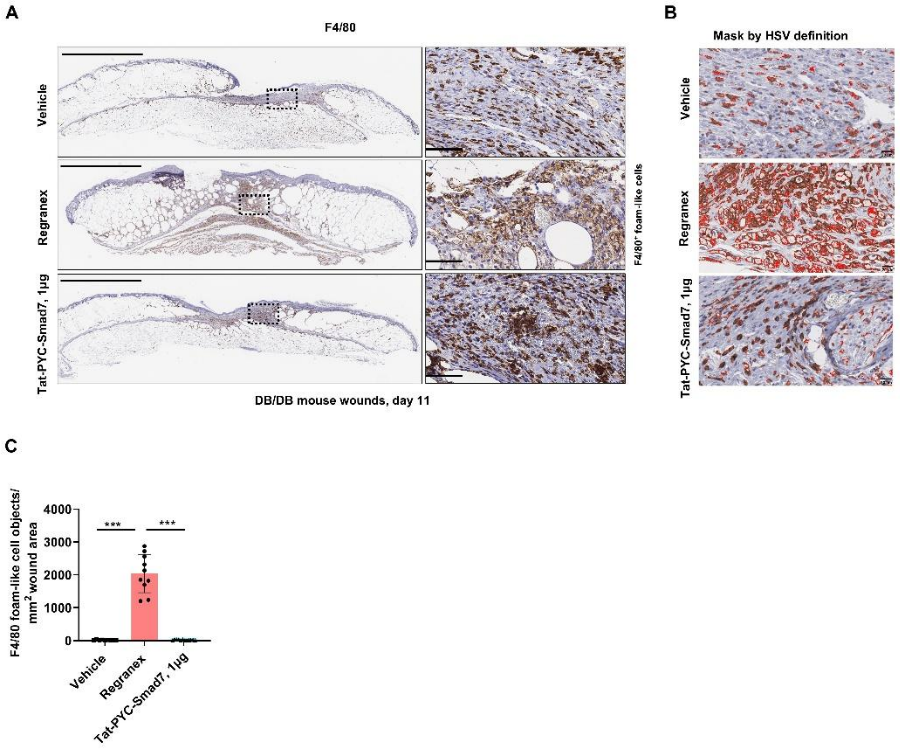


**Sup. Fig. 4. Tat-PYC-Smad7 treatment did not induce foam-like cells as Regranex.** **(A)** Representative images of wounds in db/db mice treated with vehicle, Regranex, or Tat-PYC-Smad7. Wounds were harvested on day 11 and stained for F4/80 to identify macrophages. Areas enclosed by the dotted black boxes are magnified and shown on the right. **(B)** Foam-like F4/80$^+$ cells were quantified based on hue, saturation, and value (HSV; H: 4–36, S: 35–101, V: 158–218) threshold identification and morphological characteristics using cellSens software. Red contour masks indicate identified foam-like cell objects based on HSV signal intensity, with a minimum object size threshold of 15 pixels. **(C)** Quantification of F4/80$^+$ foam-like cells per mm$^2$ wound area across treatment groups. Data are presented as mean ± SD. Statistical analysis was performed using one-way ANOVA with Tukey's multiple comparisons test. ***$P$ < 0.001. Scale bars: 1 mm (left) and 50 μm (right) in (A); 20 μm in (B). Each group included 8–10 wound samples from 4–

5 mice per experiment. Data are representative of at least three independent experiments.

HSV, hue-saturation-value; H, hue; S, saturation; V, value of intensity.

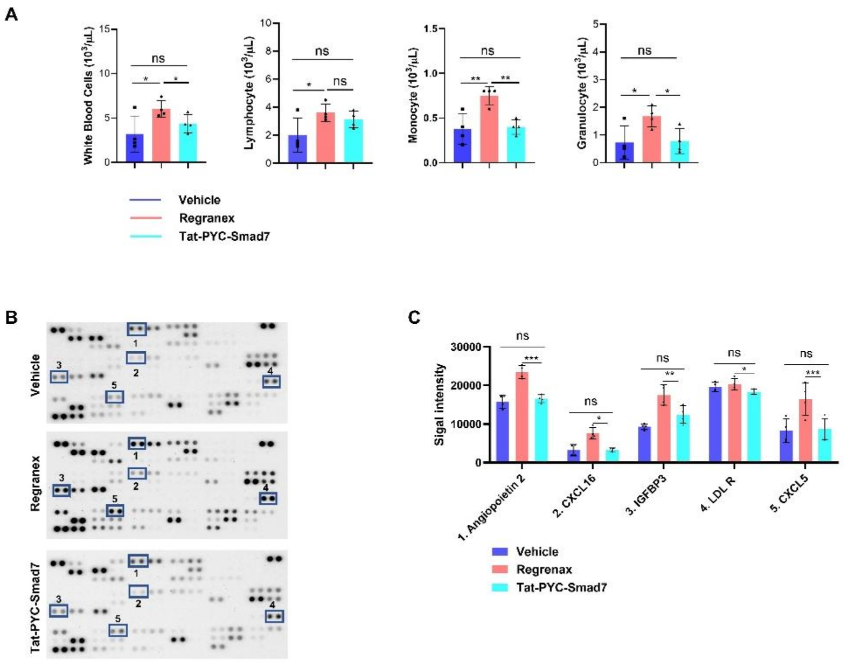

**Sup Fig.5. Wounded db/db mice treated with Tat-PYC-Smad7 did not alter blood cell counts or have significant circulating cytokine changes**. **(A)** Wounds established in db/db mice were treated with vehicle, Regranex, or Tat-PYC-Smad7. On day 11, blood was collected and analyzed by complete blood count. 200μl plasma from treated animals was applied to antibody arrays. Representative proteomic array image **(B)** and quantification of signal intensity **(C)** comparing cytokine profiles among Vehicle-treated, Regranex-treated, and Tat-PYC-Smad7-treated db/db mice. 4 blood samples in each group were quantified using unpaired t test for pairwise comparison in (A and C). Data represents the mean ± SEM. **P* <0.05, ***P* < 0.01, ****P* < 0.001, ns indicates not significantly different. LDL R, LDL receptor.

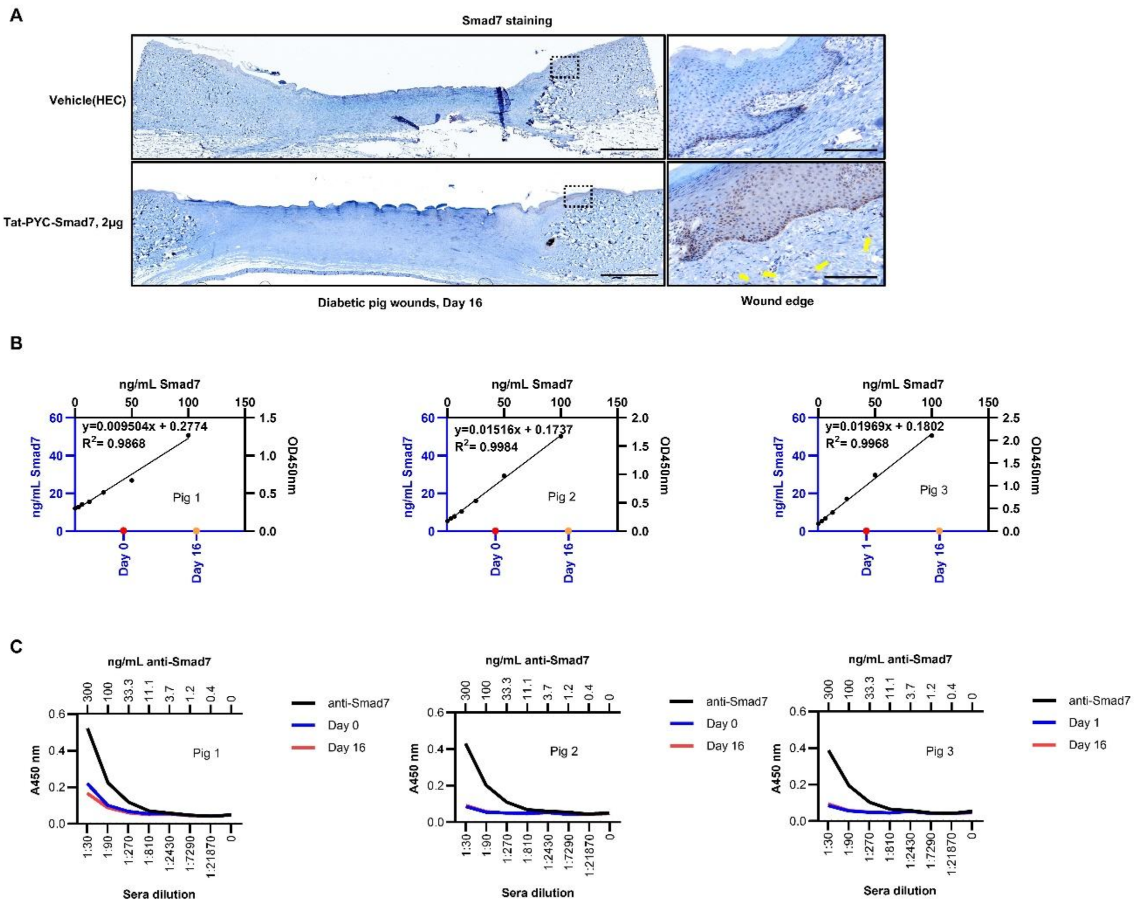


**Sup Fig.6. Tat-PYC-Smad7 penetrated to keratinocytes and stromal cells of wounded diabetic pig skin without inducing ADA**. **(A)** Immunohistochemistry staining using antibody specific to c-terminal human Smad7 revealed Tat-PYC-Smad7 within nuclei of the epidermal keratinocytes and stromal cells (yellow arrows) at the wound edge in protein-treated wounds 2 hours after application, while endogenous Smad7 was absent in the vehicle-treated wounds. Area enclosed by the dotted black box is magnified and shown on the right. Scale bars, 1mm(left), 100μm(right). **(B)** Pig serum samples were spiked with 0 to 100 ng/mL Smad7 protein to generate standard curves for quantification. The amount of Smad7 in pre- and post-treatment serum samples was measured against these standard curves. Each panel represents a different pig (Pig 1, Pig 2, and Pig 3). **(C)**

ADA ELISA was performed on diluted serum samples collected from treated pigs pre-treatment (Day 0 or Day 1) and post-treatment (Day 16) to assess the presence of anti-Smad7 antibodies. A control standard containing anti-Smad7 antibody was used for comparison. No detectable ADA response against Tat-PYC-Smad7 was observed in diabetic pigs after 16 days of wound treatment. HEC, hydroxyethyl cellulose; ADA, anti-drug antibody.

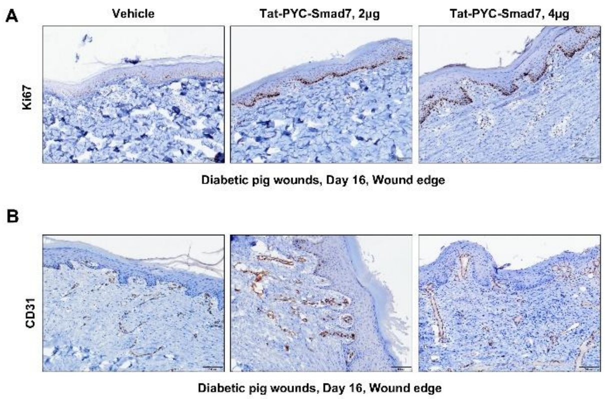


**Sup Fig.7. Tat-PYC-Smad7 promoted keratinocyte proliferation and angiogenesis in diabetic pig wounds.** Representative immunohistochemistry images and quantification of diabetic wounds from pig 2 and pig 3 stained for Ki67 **(A)** and CD31 **(B)**. Scale bars, 100µm for all images. 7-8 wound samples in each group were quantified using unpaired t test for pairwise comparison. Data represents the mean ± SEM. $^{**}P < 0.01$, $^{***}P < 0.001$.

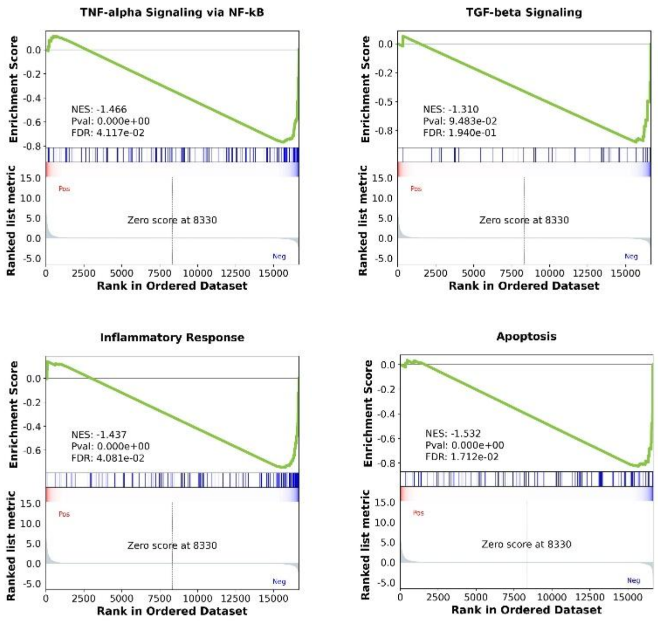


**Sup Fig.8. Tat-PYC-Smad7 inhibited TGFβ/NFκB signaling in db/db wounds.** Gene expression analysis of day 9 and day 11 wounds treated with Tat-PYC-Smad7 or vehicle were analyzed by GSEA using the Hallmark gene sets. Enrichment plots of the four indicated gene sets are presented. GSEA, gene set enrichment analysis. NES, Normalized enrichment score.

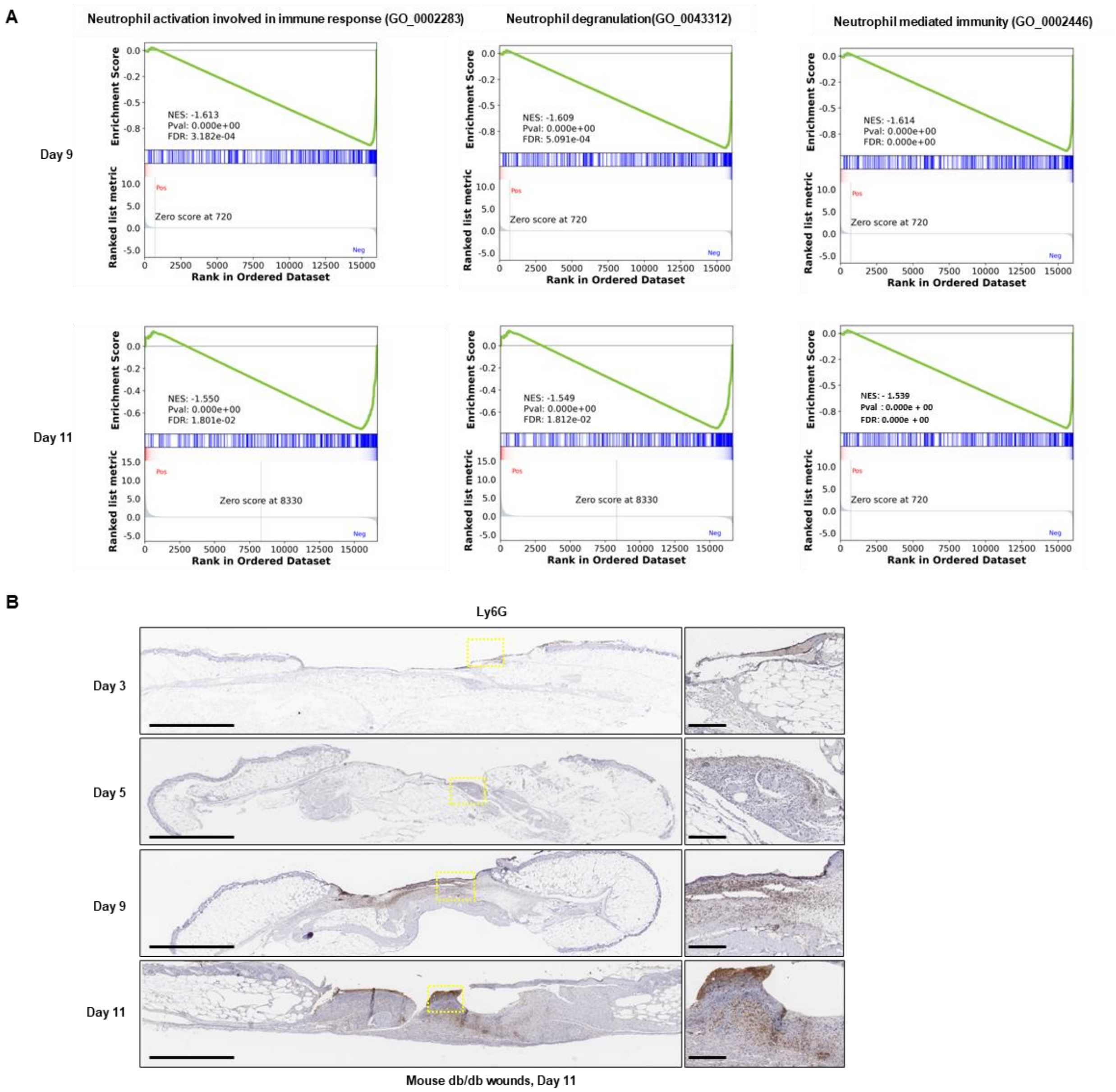


**Sup Fig.9. Tat-PYC-Smad7 blunted neutrophil activity during the chronic stage of db/db wounds. (A)** Gene expression analysis of day 9 and day 11 wounds treated with Tat-PYC-Smad7 or vehicle were analyzed by GSEA using GO biological process gene sets. Enrichment plots indicate a significant attenuation of pathways related to neutrophil activities by Tat-PYC-Smad7 treatment in db/db wounds. **(B)** Representative IHC images of Ly6G+ neutrophils in db/db wound tissues collected on days 3, 5, 9, and 11 after wounding. The areas enclosed by the dotted boxes are magnified and shown on the right. Scale bars: 1 mm (left), 100 μm (right). GSEA, gene set enrichment analysis; GO, Gene Ontology.

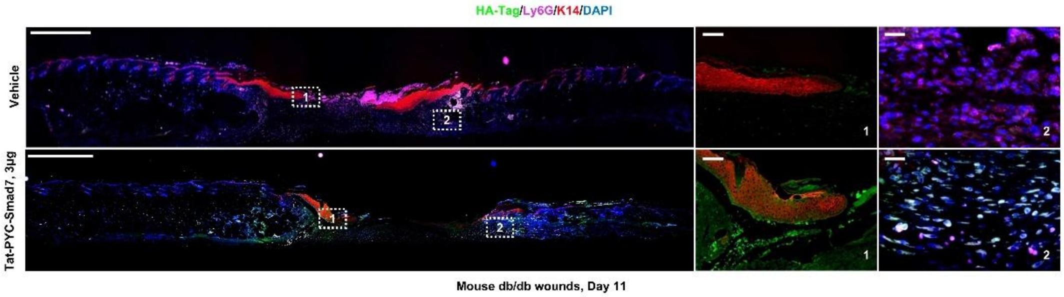


**Sup Fig. 10. Tat-PYC-Smad7 penetrated epithelial cells and Ly6G+ neutrophils in db/db wounds.** Immunofluorescent staining of HA to detect HA-tagged Tat-PYC-Smad7 near the wound in db/db mice. The areas enclosed by the dotted boxes are magnified and shown on the right highlight the presence of Tat-PYC-Smad7 in both epithelial cells and neutrophils. Scale bars: 1mm (left), 100μm (right). K14, keratin 14.

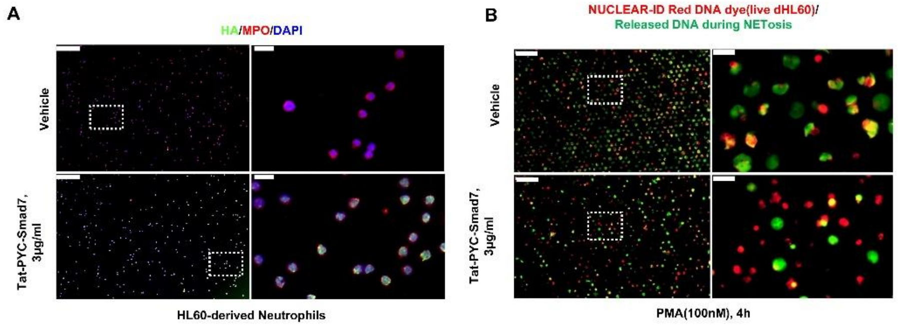


**Sup Fig. 11. Tat-PYC-Smad7 reduced PMA-induced NETosis in dHL60 cells. (A)** Representative immunofluorescence images showing HA-tagged Tat-PYC-Smad7 localization in HL60-derived neutrophils treated with Vehicle or Tat-PYC-Smad7 (3 μg/mL). Cells were stained for HA (green), MPO (red), and DAPI (blue). **(B)** Representative images of PMA-induced NETosis in dHL60 cells treated with Vehicle or Tat-PYC-Smad7 (3 μg/mL) for 4 hours. Co-localization of red and green signals was considered as objects undergoing NETosis. Representative staining areas enclosed by dotted frames in overview images are magnified on the right. Scale bars: 100 μm (left), 10 μm (right). PMA, Phorbol 12-Myristate 13-Acetate.

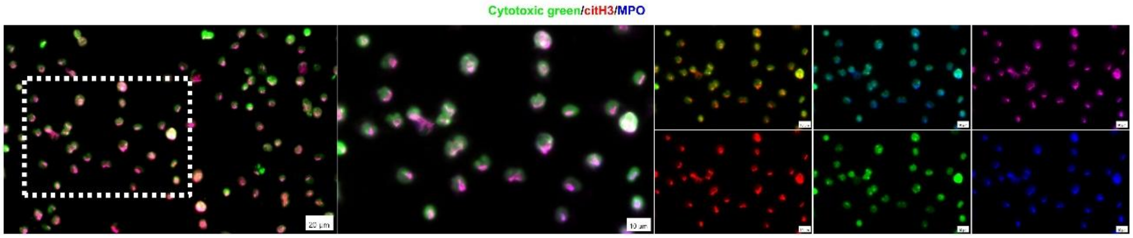


**Sup Fig. 12. NETs labeled by Cytotoxic Green colocalized with citH3-positive neutrophils and MPO.** Representative immunofluorescent images of dHL60 cells preincubated with Cytotoxic Green DNA dye and stained with MPO and citH3 antibodies 2 hours after PMA-induced NETosis. Scale bars: 20 μm for the left panel; 10 μm for the rest of the images. dHL60, differentiated HL-60; CitH3, citrullinated histone 3; MPO, myeloperoxidase.

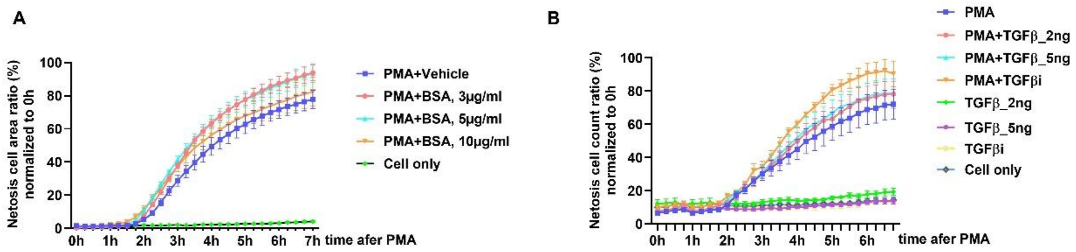


**Sup Fig.13. The effect of Tat-PYC-Smad7 on NETosis was not associated to TGFβ signaling.** Real-time quantification of PMA-induced NETosis in BSA- **(A)** or TGFβ1/TGFβi-treated **(B)** dHL60 cells using IncuCyte. Data are representative of 2 or 3 independent experiments. Data represents mean ± SEM. BSA, bovine serum albumin; TGFβi, TGFβ inhibitor (LY2109761).

A

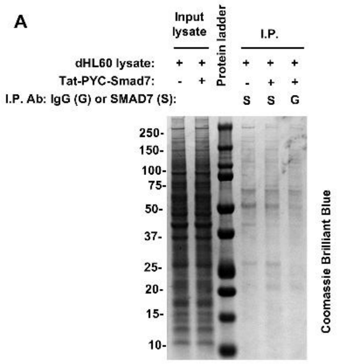


B

| *Accession Number* | *ID* | *Taxonomy* | *Vehicle + Smad7 Ab* | *Tat-PYC-Smad7 + Smad7 Ab* | *Tat-PYC-Smad7 +IgG* |
|---|---|---|---|---|---|
| P05164 | MPO | Homo sapiens | 433 | 680 | 238 |
| Q9UHX1 | PUF60 | Homo sapiens | 167 | 268 | 171 |
| P17844 | DDX5 | Homo sapiens | 117 | 328 | 146 |
| Q9H1R3 | MYLK2 | Homo sapiens | 3 | 198 | 158 |
| O00571 | DDX3X | Homo sapiens | 93 | 250 | 126 |
| Q14554 | PDIA5 | Homo sapiens | 96 | 312 | 137 |
| Q16630 | CPSF6 | Homo sapiens | 167 | 410 | 167 |
| Q15393 | SF3B3 | Homo sapiens | 71 | 171 | 108 |
| Q13435 | SF3B2 | Homo sapiens | 71 | 135 | 88 |
| O15105 | SMAD7 | Homo sapiens | 0 | 120 | 16 |

**Sup Fig.14. Tat-PYC-Smad7 binds MPO. (A)** Cell lysates from dHL60 cells treated with Tat-PYC-Smad7 or Vehicle control during PMA-induced NETosis were used for pulldown of IgG or Smad7. Coomassie Brilliant Blue staining of lysates and eluted pulldowns is presented. **(B)** Mass spectrometry spectral counts for proteins in each pull down. dHL60, HL60-differentiated cells; MPO, myeloperoxidase; I.P., immunoprecipitation.

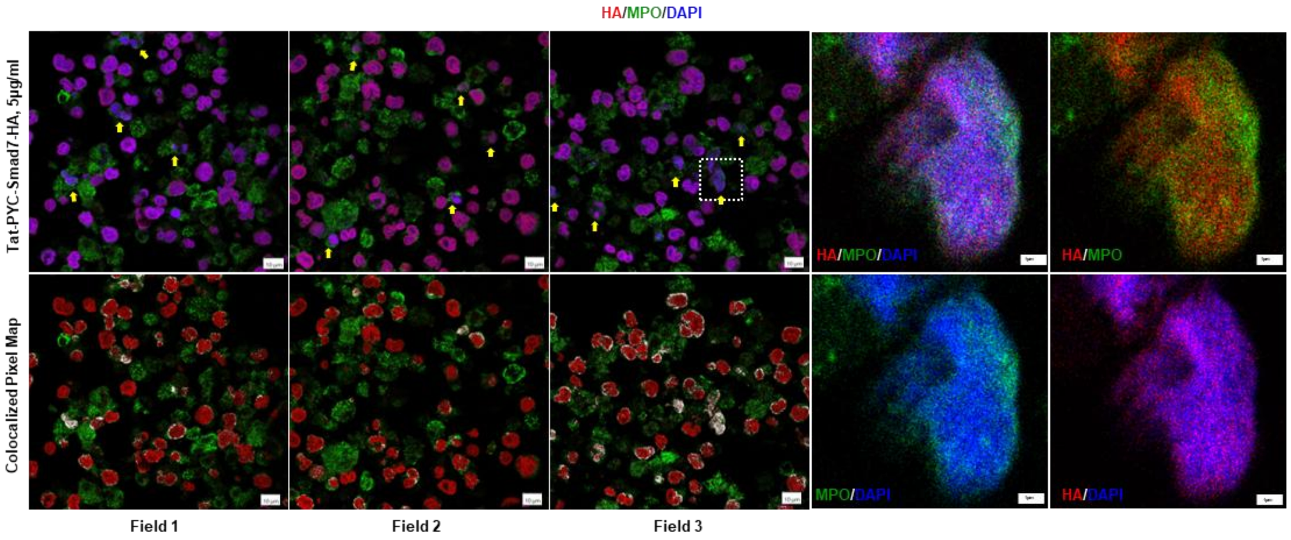


**Sup Fig. 15. Tat-PYC-Smad7 colocalization with MPO and DNA at the late stage of PMA-induced NETosis.** Representative confocal images showing nuclear colocalization of HA-tagged Tat-PYC-Smad7 (5 µg/mL) with MPO in PMA-stimulated (2.5 h) dHL60 cells. Cells were stained for HA (red), MPO (green), and nuclei (DAPI, blue). Yellow arrows indicate representative cells showing nuclear colocalization of HA and MPO. Colocalized Pixel Maps were generated using Fiji/ImageJ Colocalization Threshold plugin (bottom panels), where white pixels indicate colocalized HA and MPO signals. Enlarged images of regions indicated by dashed boxes are shown on the right. Scale bars: 10 µm (left), 1 µm (right). CitH3, citrullinated histone 3; MPO, myeloperoxidase; PMA, Phorbol 12-myristate 13-acetate; dHL60, differentiated HL-60 cells.

**Sup Table 2. Quantification of colocalization between HA and MPO signals across three different fields.**

| Metric | Field 1 | Field 2 | Field 3 | Explanation |
|---|---|---|---|---|
| Rcoloc (Thresholded Pearson's R) | 0.878 | 0.926 | 0.937 | Correlation only in regions where both HA and MPO signals are strong (above threshold). Reflects true overlap. |
| Manders M1 Thresholded | 27.69% | 22.45% | 19.63% | Percentage of strong HA signal (above threshold) overlapping with MPO (background excluded). |
| Manders M2 Thresholded | 27.55% | 23.68% | 20.23% | Percentage of strong MPO signal (above threshold) overlapping with HA. |

**Sup Table 3. Primary antibodies used in the manuscript**.

| Reagents | Manufacturer | Target Species | Catalog No. | Application |
|---|---|---|---|---|
| Smad7 Ab | Novus | Mouse, Pig | NBP1-87728 | WB, IP, IHC, MSI |
| Keratin 14 Ab | Biolegend | Mouse, Pig | 50-112-9910 | IF |
| F4/80 (D2S9R) Ab | Cell Signaling Technology | Mouse | 70076 | IHC |
| Ki67 (D3B5) Ab | Cell Signaling Technology | Mouse, Pig | 12202 | IHC, IF |
| CD31 (PECAM-1) Ab | Cell Signaling Technology | Mouse, Pig | 77699 | IHC, IF |
| HA-Tag (C29F4) Ab | Cell Signaling Technology | Mouse, Pig, Human | 3724 | ICC, IF, MSI |
| Ly6G (E6Z1T) Ab | Cell Signaling Technology | Mouse | 87048 | IHC, MSI |
| PAD4 [OTI4H5] Ab | Abcam | Human | ab128086 | WB |
| Histone H3 (citrulline R2 + R8 + R17) Ab | Abcam | Mouse, Pig, Human | ab5103 | MSI, WB, ICC |
| GAPDH (D16H11) Ab | Cell Signaling Technology | Human | 5174S | WB |
| Histone H3 Ab | Abcam | Mouse, Pig, Human | ab1791 | WB |
| Myeloperoxidase (E1E7I) Ab | Cell Signaling Technology | Human | 14569 | IP, IF, ICC, MSI |
| Myeloperoxidase Ab | Dako | Pig | GA51161-2 | MSI, IHC |
| Smad7 Ab | R&D | Pig, Human | MAB2029 | ELISA |
| αSMA Ab | Cell Signaling Technology | Pig, Mouse | 19245 | MSI, IF |
| Myeloperoxidase (E1E7I) Ab (Alexa Fluor® 647 Conjugate) | Cell Signaling Technology | Human | 17646 | IF |
| HA-Tag (6E2) Ab (Alexa Fluor® 488 Conjugate) | Cell Signaling Technology | Human | 2350 | IF |
| Lamin B1 (E6M5T) Ab | Cell Signaling Technology | Human | 17416 | MSI |

**Sup Table 3. Primary antibodies used in the manuscript**. IHC, immunohistochemistry; IF, immunofluorescence; ICC, immunocytochemistry; WB, western blot; IP, immunoprecipitation; MSI, multispectral imaging; ELISA, enzyme-linked immunosorbent assay.

**Supplementary methods**

**db/db mouse wound model**

8-12 weeks old, genetically diabetic, female BKS.Cg-Dock7m+/+Leprdb/J (db/db) mice (JAX000642) were purchased from Jackson Laboratory. Mice were acclimatized for 1 week, and their blood glucose levels were measured using a glucometer after 16 hours of fasting to confirm levels above 300 mg/dL before wound establishment. All mice were randomly grouped to control for minimal environmental variation. During the wounding procedure, mice were anesthetized with 5% inhaled isoflurane, and the dorsal surface of the mouse was shaved with an electric shaver. The surgical site was disinfected with 70% ethanol. Two pairs of dorsal full-thickness wounds were created using a 6 mm biopsy punch to excise the epidermis, dermis, and panniculus carnosus tissues. After wounding, mice were placed on a warm pad and monitored (via toe pinch) every 15 minutes until fully recovered. Liquid ibuprofen (0.2 mg/mL) was provided in drinking water continuously for 24 hours before and after the procedure. Wounds were then treated and monitored every other day. For treating wounds with Tat-PYC-Smad7, 10 µL of formulated vehicle or Tat-PYC-Smad7 gel (0.25, 0.5, and 1 µg dose) was applied evenly to each wound using a pipette equipped with filter tips. Regranex was administered using a clean cotton swab to ensure even application to the wound beds. After approximately 5 minutes, once the formulation/gel was fully absorbed, a 3M Tegaderm Transparent Film Dressing (Frame

Style, 3M™ 7 x 6 cm, 3M Healthcare, MMM1624W) was applied to completely cover the four wound beds. All treatments were administered to the mice every other day from day 1 to the endpoint of the experiment on day 3, 7, 8, 9, 11, or 25. Digital photographs of wounds were captured during treatment at a consistent distance with a calibration scale. Wound areas were calculated using a customized Python script with semi-automatic segmentation and the scikit-image library (version 0.19.3, https://doi.org/10.7717/peerj.453). Mice were euthanized using $CO_2$ asphyxiation followed by cervical dislocation. Wound tissues from identical locations were collected for further histological evaluation, lysis, or RNA extraction. Blood samples were collected in EDTA-coated tubes, and fresh anticoagulated blood samples were sent to the University of Colorado's Comparative Pathology Shared Resource Laboratory for CBC analysis. Plasma samples were collected by centrifuging blood at 1,500 rpm for 10 minutes at 4°C and stored at -80°C until further use.

**Diabetic porcine wound model and Tat-PYC-Smad7 treatment**

Diabetic porcine wound model was established by Sinclair Research Center, LLC (currently Altasciences-Columbia pre Clinical) following their approved animal protocols (Auxvasse, MO). Yucatan mini-pigs (castrated males, 27–30 kg) were used for the experiments. Diabetes was induced using an established method involving a low-dose Streptozotocin injection (50 mg/kg) [1] to achieve fasting blood glucose levels of 150 mg/dL or above at study enrollment. Animals were observed and examined by veterinary staff throughout the entire study. Prior to the wounding procedure, animals received a subcutaneous injection of buprenorphine sustained release (0.2 mg/kg) and an intramuscular (IM) injection of ketoprofen (2.2 mg/kg) as analgesic treatment. For prophylaxis, Excede® (5 mg/kg, IM) and an intravenous injection of cefazolin (20 mg/kg) were administered as antibiotics. On experimental day 1, animals were pre-anesthetized with atropine (0.04 mg/kg, IM) and a combination of tiletamine/zolazepam (Telazol) and xylazine (2.2 mg/kg, IM). Each animal was intubated endotracheally and maintained using 5% isoflurane. Twelve full-thickness 2 cm diameter excisional wounds (6 sites per side) were established along the dorsal column of the spine using a skin punch. Immediately after wounding, hemostasis was achieved by applying gauze sponges soaked in epinephrine solution (1:10,000 diluted in 0.9% sterile saline). The area around the wounds was wiped clean to ensure the peri-wound area was dry prior to treatment. For Tat-PYC-Smad7 treatment, each wound was covered with a 3M Tegaderm Transparent Film Dressing (Frame Style, 3M™ 7 x 6 cm), and the dose formulation was applied by injecting Tat-PYC-Smad7 or vehicle gel through the film. After dose application, a second layer of Tegaderm was applied to the wounds, followed by an adhesive dressing pad. The entire wound area was covered with a foam pad and vet wrap to secure the dressing materials. During dressing changes, the area surrounding the wound was moistened with sterile saline to facilitate dressing removal and cleansed with 70% alcohol. All wounds were gently irrigated with sterile saline and wiped with gauze prior to dose administration. All pigs were dosed every other day from experimental day 1 to termination on day 16. The treatment dose started at 0.5 mL per wound, adjusted to 0.25 mL per wound on day 13 as the wound size decreased, and reduced to 0.1 mL per wound on day 16. Digital photographs of the wound sites were taken before each dose administration, with every image including a colored ruler as a scale bar and appropriate animal identifiers (animal ID, site number, and study day/date of collection). Wound areas were quantified using cellSens Dimension software (Olympus, version 2.3). Animals were sacrificed two hours after the final treatment dose on post-wounding day 16. Wound samples were collected for histological evaluation. Serum samples were collected pre-treatment (Day 0 or Day 1) and post-treatment (Day 16) from blood drawn without anticoagulants. After allowing the blood to clot at room temperature for 60 minutes, the samples were centrifuged at 3,000 rpm for 10 minutes at 4°C and stored at -80°C until further use.

**ELISA to detect systemic Tat-PYC-Smad7 from local application.**

We generated a sandwich ELISA to detect Tat-PYC-Smad7 as previously described[2]. Rabbit anti-Smad7 (Novus, NBP1-87728) was coated onto 96-well plates (Nunc-Immuno, 446612) overnight and served to capture Smad7 from standards and samples. Mouse monoclonal antibody (R&D, MAB2029) that recognizes the c-terminus of human Smad7 was used for detection. We used HRP-conjugated horse anti-mouse-IgG (Cell Signaling Technologies) to detect captured immune complexes and 1-Step Ultra TMB-ELISA substrate (Thermo Fisher Scientific) to detect HRP activity, followed by acid quenching and detection on a plate reader at OD450. We spiked Tat-PYC-Smad7 into untreated pig serum to establish a standard curve and compared Smad7 levels in pig serum samples pre- (Day 0 or Day 1) and post- (Day 16) Tat-PYC-Smad7 treatment.

**Anti-Drug Antibody (ADA) ELISA**

An ADA ELISA was developed to assess anti-Smad7 antibodies in pig serum. Mouse anti-Smad7 antibody (R&D Systems, MAB2029) was used to generate a reference standard. The assay conditions were optimized by titrating secondary antibodies to minimize cross-reactivity while generating similar on-target signal using anti-mouse HRP (Cell Signaling Technology, 7076) at 1:5,000 and anti-pig HRP (Jackson ImmunoResearch, 114-035-003) at 1:25,000. 96-well microplates (Immulon 4B, Thermo Fisher Scientific, 3855) were coated with recombinant Smad7 protein (20 μg/mL in PBS) and incubated overnight at 4°C. Wells were blocked with casein-based blocking buffer (Thermo Scientific, PI37528) for 1 hour at room temperature, followed by addition of three-fold serial serum dilutions (1:30, 1:90, 1:270, 1:810, 1:2430, 1:7290, 1:21870 and 0). Bound antibodies were detected using anti-mouse HRP (1:5,000) and anti-pig HRP (1:25,000), followed by TMB substrate (Sigma, T8665) and 1M sulfuric acid stop solution. OD450 was measured using a Bio-Rad iMark Microplate Reader. ADA positivity was determined based on OD450 values exceeding background levels and aligning with the reference standard.

**Immunofluorescence (IF), Immunohistochemistry (IHC), Multispectral Imaging (MSI) and Immunocytochemistry (ICC)**

IF/IHC/MSI staining was performed on formalin-fixed paraffin-embedded (FFPE) skin sections. ICC was conducted to detect protein expression and localization in cultured cells. The IF and IHC staining procedures followed previously described protocols[3]. After primary antibody incubation, IF slides were stained with Alexa Fluor-conjugated secondary antibodies (Invitrogen™) and counterstained with DAPI. IHC slides were treated with SignalStain® Boost IHC Detection Reagent (HRP, Rabbit) (Cell Signaling Technology, catalog number 8114) or SignalStain® Boost IHC Detection Reagent (HRP, Goat) (Cell Signaling Technology, catalog number 63707), followed by visualization using the DAB Quanto Detection System (Epredia, catalog number TA-125-QHDX). For co-staining of Ly6G/CitH3/DAPI and MPO/CitH3/DAPI, MSI was performed using the Opal™ 4-Color Fluorescent IHC Kit (Akoya Biosciences, catalog number NEL810001K) according to the manufacturer's instructions. For ICC, 1 million cells were seeded into

Ibidi eight-well removable slides precoated with 0.01% poly-L-lysine solution (Sigma-Aldrich) and allowed to adhere for 20 minutes. Cells were stained using the Immunofluorescence Application Solutions Kit (Cell Signaling Technology, 12727) following the manufacturer's protocol. Primary antibodies used for IF/IHC/MSI/ICC are listed in Sup Table 3. Secondary antibodies for IF and ICC included Alexa Fluor™ 488-conjugated anti-mouse IgG (Invitrogen™, A-11001), Alexa Fluor™ 594-conjugated anti-rabbit IgG (Invitrogen™, A-11012), and Alexa Fluor™ 647-conjugated anti-rabbit IgG (Invitrogen™, A-21244). After staining, MSI/IF/ICC slides were mounted with ProLong™ Gold Antifade Mountant (Invitrogen™, P36930) and cured in the dark at room temperature for 24 hours. The slides were then stored at 4°C until imaging.

**Imaging and Quantification**

Imaging was performed using an Olympus IX83 microscope system for IF/ IHC/MSI slides, while confocal imaging was conducted using the Leica TCS SP8 STED 3X. For image quantification, sequential 10x or 20x images were captured along the basement membrane, with three images per slide. Quantification was performed using cellSens Dimension software (Olympus, version 2.3), and the average of the three images was considered the representative value for each sample slide. Nuclear Ki67-positive cells were quantified as the percentage of positive cell area relative to the epidermal area delineated by keratin 14 staining. CD31-, Ly6G/CitH3/DAPI-, and MPO/CitH3/DAPI-positive cells were quantified as the percentage of positive cell area relative to the total skin area within the wound area. The quantification of F4/80$^{+}$ foam-like cells was performed using Count&Measure function of cellSens Dimension software (Olympus, version 2.3) based on hue, saturation, and value (HSV) thresholding and morphological criteria. To specifically identify foam-like F4/80$^+$ cells, images were processed using an HSV segmentation mask with the following threshold parameters: Hue: 4–36, Saturation: 35–101, Value: 158–218. The minimum object size was set to 15 pixels. Colocalization analysis was performed using the Colocalization Threshold plugin in Fiji (ImageJ) on confocal images of HA and MPO. Automatic thresholding (Costes method) was applied to remove background noise. Thresholded Pearson's correlation coefficient (Rcoloc; range: –1 to +1) measured linear correlation within signal-positive regions above threshold, with values near +1 indicating strong colocalization. Thresholded Manders' coefficients (M1 and M2) quantified the percentage of HA or MPO signal overlapping with the other above threshold, ranging from 0% (no overlap) to 100% (complete overlap), providing a refined assessment of specific, biologically relevant colocalization.

**Trichrome Staining and Movat Pentachrome Staining**

Trichrome staining and Movat Pentachrome staining were performed using Trichrome Stain Kit (Abcam, ab150686) and Movat Pentachrome Stain Kit (Abcam, ab245884) following the manufacturer's instructions. Elastic fibers were identified as black to blue/black striae in Movat Pentachrome staining.

**RNA-Seq Data Analysis**

Tissue samples for RNA-Seq analysis were collected, rapidly frozen in liquid nitrogen, and stored at -80°C. RNA extraction was performed using an RNA Plus mini-Kit from Qiagen (Qiagen, 74136). The RNA sequencing and library preparation was conducted at the University of Colorado Denver Cancer Center's Genomics Shared Resource. NuQuant Universal Plus mRNA seq kit was used to prepare mRNA sequencing libraries. Sequencing was performed on an Illumina NovaSEQ 6000 with 2x150 paired-end reads at 80 million reads per sample depth. Quality control was conducted by FastQC (https://www.bioinformatics.babraham.ac.uk/projects/fastqc/) and FastQ Screen (https://www.bioinformatics.babraham.ac.uk/projects/fastq_screen/). The RNA-seq reads were processed with BBDuk (BBMap – Bushnell B. – sourceforge.net/projects/bbmap/) and aligned to mouse genome GRCm38.p6 (release 96) with STAR [4]. The differential expression analysis was performed on TMM (trimmed-mean M values) normalized count data using negative binomial generalized linear model and likelihood ratio test[5,6] implemented in the edgeR package (version 4.0.11, https://doi.org/10.5281/zenodo.3748085)[7]. The Benjamini–Hochberg multiplicity correction was applied on p-values to calculate false discovery rates. GSEA was performed using pre-ranked GSEA[8] and Enrichr [9] implemented by the GSEApy package (version 1.1.2, https://doi.org/10.5281/zenodo.3748085), with pathways from GO biological process, KEGG, and hallmark gene set collection from the Molecular Signatures Database. The pre-ranked GSEA utilized all identified genes pre-ranked by the negative log transformed p-values with the sign of log2 fold change. The Enrichr handled a list of differentially expressed genes with p-values less than 0.05 and absolute log2 fold changes greater than 0.6

**Proteome profiler antibody arrays**

Tissue lysates from db/db mouse wounds and plasma samples were analyzed using Proteome Profiler Antibody Arrays (R&D Systems). For tissue lysis, wound tissues were placed in RIPA buffer (Cell Signaling Technology, 9806) supplemented with Protease Inhibitor Cocktail (Roche, 4693124001). The tissues were finely cut, homogenized, and centrifuged at 10,000g for 10 minutes at 4°C to remove debris. Protein concentrations were measured using the Pierce™ BCA Protein Assay Kit (ThermoFisher, 23225) and adjusted to equal amounts before analysis. Plasma samples were collected from euthanized mice at the endpoint of the wound study as described above. Tissue lysates (500 µg) or freshly thawed mouse plasma samples (200 µL) were analyzed using the Mouse XL Cytokine Array Kit (R&D Systems, ARY028). After blocking, samples were incubated with spotted nitrocellulose membranes according to the manufacturer's instructions. Membrane HRP luminescence intensities were visualized using a ChemiDoc Imaging System (Bio-Rad), with an exposure time of 5 min, and images were saved as high-resolution TIFF files. The registered intensity of each dot was determined in duplicates by Quick Spots software (Ideal Eyes Systems), with background signals from blank control spots subtracted. Four biological replicates were measured and analyzed per condition.

**Cell culture, treatment and siRNA transfection**

HL60 cells purchased from the University of Colorado Cancer Center Cell Technologies Resource were maintained in RPMI 1640 medium (Gibco) supplemented with 10% FBS and 1% Primocin™ (InvivoGen). Cells were differentiated with culture media supplemented with 1.3% DMSO (Sigma, D2650) as previously described[10]. Differentiated HL60 cells (dHL60) were used 3 or 7 days after the initiation of differentiation. For PMA-induced NETosis, dHL60 were incubated with size exclusion chromatography (SEC) buffer (250 mM NaCl, 25 mM $NaPO_4$, pH 7.6) as vehicle control, or with Tat-PYC-Smad7 or BSA at the dose of 3, 5, and 10µg/mL (in SEC buffer) for 10min, or with recombinant human TGFβ1 (R&D, 7754-BH) at the dose of 2, 5ng/mL or TGFβ inhibitor LY2109761 at the dose of 1µg/ml for 48h, before adding 20nM or 100nM PMA to induce NETosis. For high-glucose experiments, differentiated HL60 (dHL60) cells were incubated in Ham's F-12K (Kaighn's) Medium culture medium (Gibco, 21127022) containing either normal glucose (NG) (7 mM, 1.260 g/L) or high glucose (HG) (33 mM) for 6 hours. A glucose concentration of 33 mM (equivalent to 594 mg/dL) is similar to the fed blood glucose level in BKS.Cg-Dock7m+/+Leprdb/J (db/db) mice. Mannitol (26 mM in medium with NG) was used as an osmotic control. Cells were then treated with Tat-PYC-Smad7 at concentrations of 3 and 5 μg/mL or vehicle control (SEC buffer).

For siRNA transfection, dHL60 cells were seeded at a density of 300,000 cells per well in a 6-well plate after three days of differentiation. A total of 0.02 nmol of ON-TARGETplus Human Smad7 siRNA-SMARTpool (L-020068-00-0005, Horizon Discovery/Dharmacon) or 0.02 nmol nontargeting siRNA pool (D-001810-10-05, Horizon Discovery/Dharmacon) was transfected using 1.2 μL of Lipofectamine™ RNAiMAX Transfection Reagent (Thermo Fisher, 13778100) per well. A subset of cells was harvested 48 hours post-transfection for RNA extraction and RT-PCR, as well as for cell lysis and western blotting to evaluate knockdown efficiency. The remaining cells were treated with HG or PMA to induce NETosis and monitored using IncuCyte, as described below.

**Imaging and quantification of NETosis using SIEVEWELL**

dHL60 cells were incubated for 5 minutes with the membrane-permeable NUCLEAR-ID Red DNA dye (Enzo Life Sciences, ENZ52406) to stain nuclei. Following three washes, NUCLEAR-ID Red-stained dHL60 cells were loaded into SIEVEWELL chambers (TOK, Japan). To induce and assess NETosis, RPMI medium containing 200 nM Cytotox Green Reagent (Sartorius, 4633) and 100 nM PMA (Cayman Chemical, 10008014) was added.

After 3 hours, SIEVEWELL slides were imaged using an Olympus IX83 microscope. Sequential 20x images per slide were quantified using cellSens Dimension software (Olympus, version 2.3). Co-localization of red and green signals was considered as objects undergoing NETosis.

**Live-cell NETosis observation using the IncuCyte**

For the NETosis assay, the ImageLock plate was coated with 0.1µg/mL fibronectin (Sigma, F1141) for 1 hour. dHL60 (100µL per well, 200, 000 cells) was incubated in 250nM Cytotox Green Reagent in a low fluorescence media Ham's F-12K (Kaighn's) Medium (Gibco, 21127022). Cells were scanned at the baseline level (0h). Live-cell imaging was taken every 10 or 30 minutes or 1 hour by IncuCyte® Live-Cell Analysis System (Sartorius) for at least 6 hours. The data was processed using IncuCyte® software (Sartorius).

**Cell pellet collection and embedding**

dHL60 cells were incubated with either SEC control or Tat-PYC-Smad7 (5 µg/mL) for 10 minutes, followed by treatment with 100 nM PMA to induce NETosis for 1 hour or 2.5 hours. Cell suspensions were collected into 1.5 mL microcentrifuge tubes and centrifuged at 1,500 rpm, 18°C for 5 minutes. The resulting cell pellets were fixed with 10% neutral buffered formalin (NBF) for 15 minutes, washed twice with 70% ethanol, and resuspended in fresh 70% ethanol. Pellets were then embedded in Histogel, processed into paraffin blocks, and sectioned for histological analysis.

**Elastase Detection**

The concentration of elastase in cell supernatants or tissue lysates was measured using the NETosis Assay Kit (Abcam, ab235979) to detect the elastase production. dHL60 cells were incubated with either SEC buffer or Tat-Smad7 protein (10 μg/mL) for 10 minutes before being treated with PMA (100 nM) for 4 hours to induced NETosis. After washing, the cells were treated with S7 Nuclease for 15 minutes. Supernatants from each well were collected and assayed for neutrophil elastase. Prepared cell supernatants or tissue extracts from wounds (1 μg/mL) were incubated for 2 hours at 37°C in 200 μL of 0.1 M HEPES buffer (pH 7.5) containing 0.5 M NaCl, 10% DMSO, and 0.1 mM elastase substrate. A standard curve for substrate degradation was generated using neutrophil elastase, ranging from 0 to 36 mU per well. Substrate degradation was quantified by measuring absorbance at 405 nm.

**Quantitative Real-Time PCR**

Total RNA was extracted from cells or tissue samples using the RNeasy Plus Mini Kit (Qiagen, Cat. No. 74134). cDNA was synthesized from 0.02–2 μg of total RNA using the High-Capacity cDNA Reverse Transcription Kit (Applied Biosystems, 4368814). Relative gene expression levels were determined using TaqMan™ Gene Expression Assay (FAM) (Thermo Fisher Scientific, 4331182) with TaqMan™ Fast Advanced Master Mix (Applied Biosystems, 4444557), and reactions were performed on a QuantStudio 5 Real-Time PCR System. The TaqMan Gene Expression Assays used in this study included Human SMAD7 (Assay ID: Hs00998193_m1) and Human GAPDH (Assay ID: Hs02758991_g1). The comparative Ct (ΔΔCt) method was applied to calculate relative fold changes in gene expression, with GAPDH serving as the endogenous control.

**Western Blotting**

Protein samples were harvested with RIPA buffer (Cell Signaling Technology, 9806) supplemented with a protease inhibitor cocktail (Roche, 4693124001). Western blotting was performed using standard protocols as previously described[11] and detected using a ChemiDoc Imaging System (Bio-Rad). Antibodies used in western blotting are listed in Sup Table 3. The grayscale intensity of protein bands was quantified using ImageJ software. Relative expression levels were determined by normalizing the intensity of each target protein band to its corresponding loading control band. These normalized values were then adjusted relative to the highest intensity band, which was set to 1.

**Immunoprecipitation**

dHL60 cells were incubated with 10 μg/mL Tat-PYC-Smad7 protein or SEC buffer for 10 minutes before adding PMA (20 nM) to induce NETosis. Cells were lysed in cell lysis buffer (Cell Signaling Technology, 9803) supplemented with freshly added protease inhibitors (Roche, 4693124001). For immunoprecipitation, 1 μg of anti-Smad7 antibody (recognizing C-terminal Smad7, Novus, NBP1-87728) or 1 μg of rabbit polyclonal IgG (Novus, NB810-56910) was added to 500 μg of whole-cell lysate. Immunoprecipitation was performed by incubating the mixture with 20 μL of prewashed protein A magnetic beads (Cell Signaling Technology, 73778), followed by five washes with 500 μL of 1X cell lysis buffer (Cell Signaling Technology, 9803). The pellet was resuspended in 20-40 μL of 3X SDS sample buffer, vortexed briefly, and centrifuged. The sample was then heated to 95-100°C for 5 minutes. Beads were separated using a magnetic rack, and the supernatant was transferred to a new tube. An aliquot of 15–35 μL was loaded onto an SDS-PAGE gel for protein separation, followed by transfer to a nitrocellulose membrane for western blot analysis. Immunoprecipitated samples were also submitted to the Mass Spectrometry Proteomics Shared Resource Facility for proteomic analysis.

**Mass Spectrometry Proteomics**

**Sample preparation**

Samples were loaded onto a 1.5 mm thick NuPAGE Bis-Tris 4−12% gradient gel (Invitrogen). The BenchMark™ Protein Ladder (Invitrogen) was used as a protein molecular mass marker. The electrophoretic run was performed by using MES SDS running buffer, in an X-Cell II mini gel system (Invitrogen) at 200 V, 120 mA, 25 W per gel for 30 minutes. The gel was stained using SimplyBlue™ SafeStain (Invitrogen, Carlsbad, CA) stain and de-stained with water according to the manufacturer's protocol. Each lane of the gel was divided into 3 equal-sized bands, and proteins in the gel were digested as follows. The gel pieces were destained in 200 μL of 25 mM ammonium bicarbonate in 50 % v/v acetonitrile for 15 min and washed with 200 μL of 50% (v/v) acetonitrile. Disulfide bonds in proteins were reduced by incubation in 10 mM dithiothreitol (DTT) at 60 °C for 30 min and cysteine residues were alkylated with 20 mM iodoacetamide (IAA) in the dark at room temperature for 45 min. Gel pieces were subsequently washed with 100 μL of distilled water followed by addition of 100 mL of acetonitrile and dried on SpeedVac (Savant ThermoFisher). Then 100 ng of trypsin was added to each sample and allowed to rehydrate the gel plugs at 4 °C for 45 min and then incubated at 37 °C overnight. The tryptic mixtures were acidified with formic acid up to a final concentration of 1%. Peptides were extracted two times from the gel plugs using 1% formic acid in 50% acetonitrile. The collected extractions were pooled with the initial digestion supernatant and dried on SpeedVac (Savant ThermoFisher). Samples were desalted on Thermo Scientific Pierce

C18 Tip.

**Mass spectrometry analysis**

A 20 µL of each sample was loaded onto individual Evotips for desalting and then washed with 20 µL 0.1% FA followed by the addition of 100 µL storage solvent (0.1% FA) to keep the Evotips wet until analysis. The Evosep One system (Evosep, Odense, Denmark) was used to separate peptides on a Pepsep column, (150 um inter diameter, 15 cm) packed with ReproSil C18 1.9 um, 120A resin. The system was coupled to the timsTOF Pro mass spectrometer (Bruker Daltonics, Bremen, Germany) via the nano-electrospray ion source (Captive Spray, Bruker Daltonics).

The mass spectrometer was operated in PASEF mode. The ramp time was set to 100 ms and 10 PASEF MS/MS scans per topN acquisition cycle were acquired. MS and MS/MS spectra were recorded from *m/z* 100 to 1700. The ion mobility was scanned from 0.7 to 1.50 $Vs/cm^2$. Precursors for data-dependent acquisition were isolated within ± 1 Th and fragmented with an ion mobility-dependent collision energy, which was linearly increased from 20 to 59 eV in positive mode. Low-abundance precursor ions with an intensity above a threshold of 500 counts but below a target value of 20000 counts were repeatedly scheduled and otherwise dynamically excluded for 0.4 min.

**Database Searching and Protein Identification**

MS/MS spectra were extracted from raw data files and converted into .mgf files using MS Convert (ProteoWizard, Ver. 3.0). Peptide spectral matching was performed with Mascot (Ver. 2.5) against the Uniprot human database. Mass tolerances were +/- 15 ppm for parent ions, and +/- 0.04 Da for fragment ions. Trypsin specificity was used, allowing for 1 missed cleavage. Met oxidation, protein N-terminal acetylation, peptide N-terminal pyroglutamic acid formation set as variable modifications with Cys carbamidomethylation set as a fixed modification.

Scaffold (version 4.9, Proteome Software, Portland, OR, USA) was used to validate MS/MS based peptide and protein identifications. Peptide identifications were accepted if they could be established at greater than 95.0% probability as specified by the Peptide Prophet algorithm. Protein identifications were accepted if they could be established at greater than 99.0% probability and contained at least two identified unique peptides.

**MPO activity assay**

The EnzChek MPO Activity Assay Kit (E33856, Invitrogen) was used for the rapid and sensitive determination of MPO chlorination activity and peroxidation activity using cell lysates from dHL60 cells treated with Tat-PYC-Smad7 or Vehicle control during PMA-induced NETosis. The fluorescence intensity of each sample was measured with a fluorescence microplate reader using excitation at 485 nm and emission at 530 nm for chlorination activity, and excitation at 530 nm and emission at 590 nm for peroxidation activity. The background fluorescence measured for each zero-MPO control reaction was subtracted from each fluorescence measurement before plotting.

**Statistical Analysis.**

Statistical analysis was performed using GraphPad Prism 10 software following the steps outlined below. The ROUT method (Q = 1%) was applied to identify outliers within the group data sets. After outlier removal, the Shapiro-Wilk normality test was conducted to assess the normality of the data. For data sets following a Gaussian distribution, a two-tailed unpaired t-test was used for pairwise comparisons between two groups. For data sets that did not meet the criteria for Gaussian distribution, the Mann-Whitney test was employed for pairwise comparisons.

Multiple comparisons were conducted using either two-way ANOVA followed by Tukey’s multiple comparison test, Sadik’s multiple comparison test, or Fisher’s LSD test, or one-way ANOVA followed by Tukey’s multiple comparison test. Contingency data were analyzed using Fisher's exact test. Data were presented as the mean ± SD or the mean ± SEM. Results represent either pooled data from 2-3 independent experiments with 6–15 samples or animal subjects per group, or data from at least 3 independent experiments with 3–5 samples or 4–5 animal subjects per group.
